# From gonadogenesis to testicular cancer: Unravelling the novel regulators and identification of drug candidates against FGF9 and PTGDS

**DOI:** 10.1101/2023.06.09.544377

**Authors:** Cash Kumar, Vinay Kumar Singh, Jagat Kumar Roy

## Abstract

Sex determination is the preliminary step toward gonadogenesis in mammals. Antagonistic interactions of key regulators have been only fragmentarily mentioned so far. Therefore, exploring regulators underlying the phenomena is required to solve questions, especially regarding female gonad development and gonadal disorders in congenital or adults. Inhibiting discrepancies in PPI pathways and combating related disorders are of urgent necessity, for which novel drugs are constantly required. Here, we performed *in silico* analysis using robust bioinformatics methods, which is unprecedented work in sex determination studies, providing large-scale analysis without exorbitant wet lab procedures. Analyzed regulators were overlapped with our RNA-seq data for authentication, to obtain differentially expressed elements. Additionally, CADD approach was used to discover inhibitors for FGF9 and PTGDS to search for potential drugs combating gonadal disorders in adults. Along with druggable properties, only FGF9 and PTGDS had full-length protein structures available, among 25 key genes under investigation. Our large-scale analysis of PPIN, produced highly interacting hub-bottleneck nodes as novel genes. Further, functional enrichment analysis revealed importance of these regulators in gonadogenesis. We identified sex-specific novel genes, miRNAs-target pairs, and lncRNAs-target pairs, which appear to play an important role in regulation of ovary development. CADD with molecular docking, MD simulations, and molecular mechanics confirmed stability of two novel compounds, DB12884 and DB12412 that could potentially inhibit FGF9 and PTGDS respectively. Taken together our study provides valuable information regarding involvement of crucial regulators in antagonistic mechanism of gonadogenesis and their related disorders, which will further assist in refining wet lab experiments.

## 1. Introduction

Sex determination and development of bipotential gonads into sex-specific gonads, testis, or ovaries is one of the fascinating phenomena during embryonic development in mammals. Development of an organ requires a well-defined cross-talk between various signaling pathways. Development of either testis or ovaries involves a sex-specific molecular pathway (Windley & Wilhem, 2015) along with antagonistic activity of key players, mainly autosomal in origin. Differential expression of autosomal genes with synergistic effect of specific regulators is the main defining factor in differentiation of male/female gonad (Eicher et al., 1988). Expression of transcription factor, Sry regulates key downstream regulator *Sox9*, which further activates *Fgf9*, and both together help in suppression of female-biased gene Wnt4. This combinatorial antagonistic signaling helps in establishment of testis-specific anatomy. However, in absence of a master regulator as *Sry*, females trigger a different set of genes i.e., *Rspo1, Foxl2, Fst, and Wnt4* known to function as main regulators of female gonad development and help in stabilization and translocation of B-catenin into nucleus. Gonadogenesis involves various molecular pathways, including WNT signaling, TGFβ including BMPs, hedgehog, Hippo signaling, and NOTCH signaling pathways. Various genome-wide studies have been conducted to understand how sex determination and gonadal differentiation are controlled at genetic level and how they are antagonistically regulated. Delicate imbalances between antagonistic mechanisms of gene regulations involved in sex determination mainly lead to congenital disorders of sexual development (DSD). Extensive research has been done on masculine pathways to understand mostly unknown area of DSDs. But knowledge gap and lack of understanding male and female antagonistic pathways mainly restricts longstanding goals of comprehending DSDs.

The major understanding of proteins involved in sex determination and DSDs revolves around approaches to identifying gene regulatory pathways, novel genes, and their associations with various biological processes. This can provide novel ideas for apprehending the molecular mechanisms and identifying therapeutic targets associated with sexual disorders. Transcriptomic studies have added enormous data to enhance knowledge of gonad development but proteomic analysis of developing gonad have not been fully elucidated. Wilhelm et al., 2005 conducted a proteomic study to analyze molecular events during gonadal development and identified proteins involved in cascade utilizing different robust proteomic approaches. Later in 2009, Evens et al., conducted another proteomic analysis with involvement of bioinformatic approaches and revealed embryonic gonadal proteins along with their co-expression at various locations, involved in a variety of molecular functions and biological processes. Since then, none of studies have provided any advancement in proteomic studies. To advance the fragmentary knowledge of overall regulatory mechanism revolving around central dogma which is essential for varying physiological functions. Regulatory mechanism not only includes coding genes but also non-coding RNAs including miRNA and lncRNA are promising regulators, controlling expression of several protein-coding genes with interaction of mRNA. However, very little is known about regulation of genes through these ncRNA in sex determination phenomena. Therefore, it is crucial to determine participation of these regulatory elements during gonad development. Previous studies did not provide any detailed knowledge of overall interaction between smRNA and their targets or between different small RNAs in that specific tissue or globally. According to NGS-based studies, testis expresses much more differentially expressed lncRNA than in any other tissue (Necsulea et al., 2014; Washietl et al., 2014; Sun et al., 2013; Chen et al., 2018). Our exhaustive literature survey confirmed that none of approaches utilized robust bioinformatics methods to find novel regulators involved in both testicular or ovarian development and their related disorders. Most of studies only provide information about interactions within specific tissue types. However, in this study, we have tried to fetch global interactions of these regulators in all tissue types providing an overview of functional interactome of key regulators. Nevertheless, studies regarding DSDs, mostly revolve around understanding mechanism in congenital stages only. It is noteworthy that, most of key regulators are well known to cause disorders related to gonads in adults also. However, therapeutic intervention still lacks limelight.

We have selected twenty-five key genes, critically involved in gonadogenesis. Selected genes are differentially expressed in male and female embryonic gonads. STRING database was used to fetch all interacting partners and PPI network was established. Further visualization and analysis were done through Cytoscape tool (Shanon et al., 2003). Topological and cluster analysis revealed hub and bottleneck nodes, which are important and highly interacting proteins of network. Further, functional enrichment analysis provided new insights into overall mechanism. Then we successively conducted miRNA-target analysis utilizing TransMir and mirTarBase, which can be analyzed to unveil TFs for activation of miRNAs genes and suppressing effect of miRNA during antagonistic mechanism of sex determination. In addition to this, we have also predicted a few long non-coding RNA and their targets from an online tool LncRRIsearch. Since *in vitro* and *in vivo* analysis of vast collection of compounds targeting a protein was tedious and expensive, therefore computer-aided drug identification has been used as an efficient alternative. PPI has emerged as a promising target for rational drug design owing to its involvement in cellular functions. Analyzing vast networks of protein-protein interactions contributes immensely to enhancing knowledge of complex biological systems (Murakami et al., 2017).Taking our study to next level, we have tried to predict drug molecules for gonadal disorders in adults using robust bioinformatics approaches. To predict new modulators of selected proteins, Fgf9 and PTGDS, we carried out virtual screening against FDA-approved compounds from drug bank. ADME and toxicity properties of obtained screened compounds were analyzed to examine drug-like properties. Binding modes of selected candidate compounds obtained from docking were studied by visual inspection and their binding stabilities were further evaluated by performing molecular dynamics simulation via Schrödinger DESMOND.

## 2. Materials and methods

### 2.1 Selection of candidate genes associated with sex determination and gonadogenesis

Candidate genes associated with sex determination and gonadogenesis were collected from literature. Altogether 25 genes (*Cbx2, Lhx9, Wt1, Nr5a1*, *Nr0b1, Emx2, Sry, Gata4, Fgf9, Sox9, Sox8, Ptgds, Amh, Fgfr1, Fog2, Dmrt1, Map3k1, Mp3k4, Dhh, Ctnnb1, Foxl2, Wnt4, Rspo1, Fst,* and *Bmp2*) were analyzed and selected (Table 4.1). These include genes expressed in bi-potential gonads viz. *Cbx2, Lhx9, Wt1, Nr5a1, Nr0b1, and Emx2*; other genes like, *Ctnnb1, Foxl2, Wnt4, Rspo1, Fst,* and *Bmp2* are ovary-specific genes and rest are testis-specific genes. Outline of study is depicted in Figure 1.

**Figure 1.**
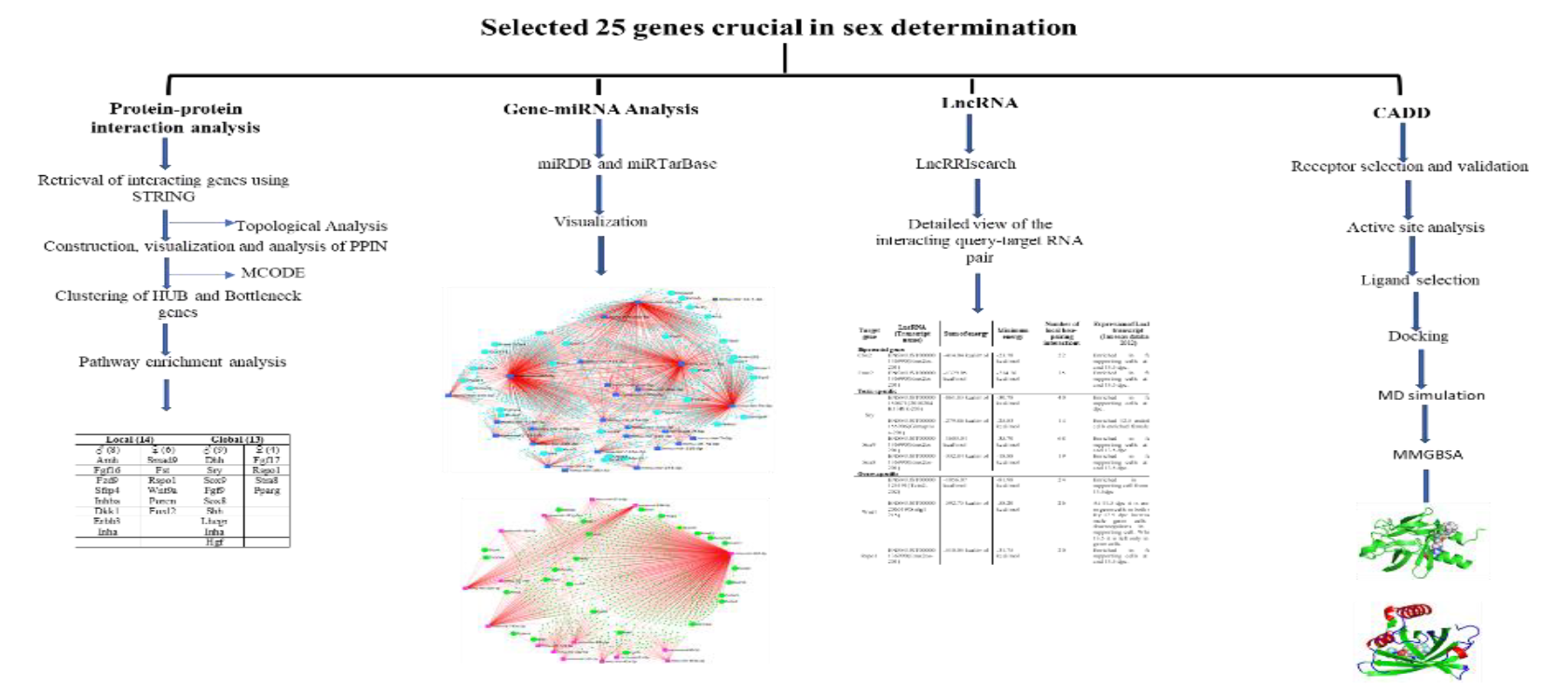
Summary of study and methodology used. Flow diagram of four types of analysis done using a set of 25 genes.

### 2.2 Construction, topological and cluster analysis of Protein-protein interaction network

The STRING database search tool (version 11.5) was used for retrieval of interacting genes/proteins. It searches protein-protein interactions, including both direct physical interaction and indirect functional correlation between proteins. System uses a scoring mechanism to give a certain weight to results and finally gives a comprehensive score (WodakPu, Vlasblom & Seraphin, 2009). To construct PPI network selected genes were analyzed at a stringency level of 0.400 to fetch out all possible interactions of each gene individually. Other parameters set, consisted of full STRING network type, selecting all options for active interaction sources with a maximum number of interactions set to 50. STRING network (.sif file) was exported to Cytoscape application, version 3.9. In procedure of protein interaction networks (PINs) analysis, Cytoscape and its in-build Network Analyzer plugin were used to calculate basic parameters ofPINs, such as degree distribution, clustering coefficient, betweenness centrality, closeness centrality, path distribution, and topological coefficients distribution.

To analyze properties of nodes and edges in network, topological analysis evaluating degree, betweenness and measures of closeness centrality was adopted. To assess topological properties, CytoHubba plugin app in Cytoscape was used. Topological measures were analysed for detecting hub and bottlenecks of network. The topological analysis yielded highly interacting genes i.e. genes that are acting both as hub and bottleneck (key genes). Using key genes, we created new cluster network along with their corresponding edges. To apprehend intersected clusters from brand-new PPI community we exploited Cytoscape plugin Molecular Complex Detection plugin (MCODE).module (cluster) figuring out extremities have been reported, together with Kappa rating (K-core)constant to five, Degree Cutoff constant to two, Max. Depth constant to 100, and Node-rating Cutoff constant to 0.2 constraining cluster length for co-expressing networks (Bader and Hogue, 2003).

### 2.3 Functional annotation and pathway enrichment analysis of key nodes

The functional annotations including Gene Ontology and KEGG pathways of genes which was performed using DAVID. Molecular function, biological processes and cellular compartments under GO were visualized and specifically looked for terms that is related to gonad development. This analysis was conducted for all genes obtained from PPIN analysis and as miRNAs mRNA targets.

### 2.4 Retrieval and construction of gene-miRNA network

MiRNA-gene network prediction was conducted to identify miRNA-regulated genes under study. miRDB (http://mirdb.org/; Wong and Wang 2015)was used to fetch out interacting miRNA of a particular target above target score of 80.Then this interaction was visualized through Cytoscape using default layout. In addition to this analysis, differentially expressed miRNAs (DEMs) from above interaction were picked using our small-RNAseq data. Utilizing miRTarBase (http://miRTarBase.cuhk.edu.cn/) we searched targets of these miRNA. From these, only differentially expressed targets were marked and visualized on mirTarBase using Forced atlas layout.

### 2.5 Prediction of differentially expressed lncRNA and their target genes

For prediction of interaction between 25 selected genes and associated lncRNA that might regulate sex differentiation pathways, we employed LncRRIsearch database (Fukunaga et al., 2019). LncRRIsearch is a web server for prediction of human and mouse lncRNA-lncRNA and lncRNA-mRNA interaction. Prediction was conducted using RIblast, which is an accurate RNA-RNA interaction prediction tool was performed by selecting a query RNA and a target RNA to get pre-calculated prediction results. To obtain a list of lncRNA for a specific gene of interest, we selected only target mRNA checkbox and inserted specific gene symbol. Based on the threshold interaction energy set to −16 kcal/mol, a list of top 100 lncRNA based on minimum and sum energy was obtained. Using transcript name of lncRNA in list we searched for differentially expressed lncRNA from Jameson database (Jameson et al, 2012) targeting our selected genes.

### 2.6 Gene-disease Association, analysis of targetable proteins

To illustrate essential protein-chemical interactions, Search Tool for Interacting Chemicals (STITCH) database (http://stitch.embl.de) was used to construct protein-chemical interaction networks.25 genes were uploaded and network was retrieved at stringency level of 0.700. All25 selected genes were reviewed for gonad-associated disorders for both embryo and adult stages and arranged in tabulated format (Table 1). Exhaustive literature mining was done and various disease databases were used for preparation of Gene-disease association table. With the aim of designing a drug for these disorders, we have checked the availability of full-length protein structure using <u>UniProt</u>, as availability of full-length protein structure is crucial for defining various domains and complete active site residues for docking calculation and drug designing purposes.

### 2.7 Druggability test

To select as a drug targets for testicular and ovarian disorders all 25 genes were tested for druggability using DragomeAI (Raies et al., 2022) and Pockdrug (http://pockdrug.rpbs.univ-paris-diderot.fr.) tools.

### 2.8 Receptor Selection and retrieval of3D structure of protein

Of 25 genes regulating sex determination involved in this study, FGF9 and PTGDS have been selected to predict drugs to combat disorders related to reproductive. For *in silico* analysis, three-dimensional (3D) crystal structure of FGF9 (PDB ID: 1IHK) and PTGDS (PDB ID: 4oru) was downloaded from RCSB-PDB (UniProt Consortium 2021: Berman et al., 2000).crystal structure of FGF9 has been deposited by Plotnikov et al. 2001 (PDB DOI: 10.2210/pdb1IHK/pdb; A: 174). And structure of PTGDS, deposited by Perduca et al., 2014 (PDB DOI: 10.2210/pdb4ORU/pdb; A: 190). These structures were further refined using MoDrefiner (Dong and Yang et al., 2011).

### 2.9 Structure Validation of both protein structure

The stereochemical stability of structure was further validated using various protein quality-based parameters. These parameters include percentage of residues in priority and allow regions, number of glycine and proline residues, and dihedral orientation, including phi (φ) and psi (ψ). PROCHECK module of PDBSum server (Laskowski et al., 2005) verifies the backbone conformation. The server VERIFY3D (Eisenberg et al., 1997) was used to check compatibility of atomic models (3D) with their primary amino acid sequences. Quality was validated using ERRAT score values based on statistics on unbounded atom interactions and atom distribution (Colovos and Yeates, 1993).

### 2.10 Active Site Analysis

The active site and binding sites prediction of protein model was analyzed using POCASA server with default parameters (Yu et al., 2010). This online tool consists of four adjustable parameters grid size, probe radius, SPF, and PDF threshold for active site prediction. Default Parameters in POCASA of Grid Size = 1Å, Probe Radius = 2Å, SPF = 16, PDF = 18 were used for calculation. Visualization of predicted active sites and their residues was done using BIOVIA Discovery Studio 2019.

### 2.11 Ligand Selection

Compounds activity checked with testicular tumorigenesis, male infertility and ovarian follicular cysts treatment were searched and retrieved. Compounds/ligands were retrieved from Drugbank database. The drug compounds selected belong to all enlisted categories of FDA approved drugs i.e. approved, experimental, investigational, nutraceutical, illicit and withdrawn.

### 2.12 Docking Calculation

Molecular docking of selected Compounds/ligands was performed using different tools viz. LibDock, CDOCKER, and Molecular dynamics simulation (Wu et al., 2003, Diller et al., 2001, Diller et al., 2003, Rao et al., 2007). Libdock is a rigid-based docking program. It calculates hotspots for selected protein with a grid positioned into binding site using polar and nonpolar probes. All compounds were ranked according to their Libdock score. CDOCKER module of Discovery Studio with implementation of a CHARMm-based docking tool was used for docking process. CHARMm energy and interaction energy, which indicates ligand-binding affinity, were calculated. CHARMm forcefield was used for receptors and ligands. During docking process, ligands were allowed to bind to residues within binding site spheres. Poses are sorted by CHARMm energy and top-scoring poses are retained.

### 2.13 Standard Dynamics Cascade – Parameters for MD simulation

The docked complexes were considered for further molecular dynamics (MD) runs.MD simulations were carried out using DESMOND simulation package included in Schrödinger Suite. Before MD simulations, Desmond default six-stage system relaxation methods were followed for equilibration. System was neutralized by adding 0.15 M salt concentrations to make up desired counter ions. 100ns MD simulations were performed for protein-ligand complex with a recording interval of 100ps.OPLS_2005 force field parameters were used in all simulations. After MD simulation, trajectories were further examined to understand ligand stability and protein conformational changes using parameters like root mean square deviation (RMSD), root mean square fluctuation (RMSF), solvent accessible surface area (SASA), radius of gyration (Rg) and molecular interactions between protein and ligand complexes. To estimate relative free energy of ligand binding, Molecular Mechanics with Generalized Bonn Surface Area (MM-GBSA) were computed with utilization of Prime using last 10 ns simulation trajectories.

## 3. RESULTS

### 3.1 Protein-protein interaction network

The Protein-Protein Interaction network of selected genes related to mammalian sex determination and gonadal differentiation, generated by STRING database was further processed, visualized, and analyzed through Cytoscape (Fig 2[A]). It consisted of 810 nodes and 12377 edges and was designated as an undirected graph. Structural properties of network were analyzed to better understand functional organization of network. Average connected component in PPI network was found to be 1, indicating that majority of proteins in network are highly interacting and suggest stronger network connectivity. Further, degree distribution in this network reflects a biological network and thereby proteins in network might efficiently communicate biological information related to gonadogenesis. In addition, PPI network has a characteristics average path length value of 2.858.clustering coefficient of this scale-free network is 0.700, which significantly describes that internal structure of this network is highly interactive and forms clusters.

### 3.2 Topological Analysis of PPI Networks for Hubs and Bottlenecks

To explore biologically important nodes in a network we utilized CytoHubba, a Cytoscape plugin, which provides 11 topological analysis methods encompassing centralities measures like degree, betweenness, and closeness. Degree measures provide hubs, betweenness highlights bottlenecks of network while closeness centrality measure shortest paths which influence most in network. These measures are divided into two categories, local-based on calculation score of a node with its direct neighbor, and global-based measuring relationship between node and whole network.

Venn diagram shows unique and common nodes yielded by local method viz. Degree, DMNC, MNC, and MCC. A total of 123 nodes are found to be common in local analysis (Figure 2 [B]). The highest number of edges which depicts interaction between nodes in network is represented by MCC measure i.e., 5485. Other measures also show a high number of edges interacting in network. Through global analysis also, high-scored edges can be obtained (Fig2[C]). After topological analysis of entire network, we finally retrieved 123 nodes yielded by local methods which consists of 4 of our input genes and 68 nodes by global measures with 13 genes in common with input data that are differentially expressed in male and female gonads. Then overlapped local and global nodes with our RNA-seq data (GEO: SUB12930291) which provide differentially expressed genes. This analysis yielded 14 genes from local method that are common from differential expression with 8 genes upregulated in males and 6 in females. On other hand, out of 68 genes of global analysis 13 genes are differentially expressed. In which 9 genes are upregulated in males and 4 genes in females (Figure 3; Table 2).

**Figure 2.**
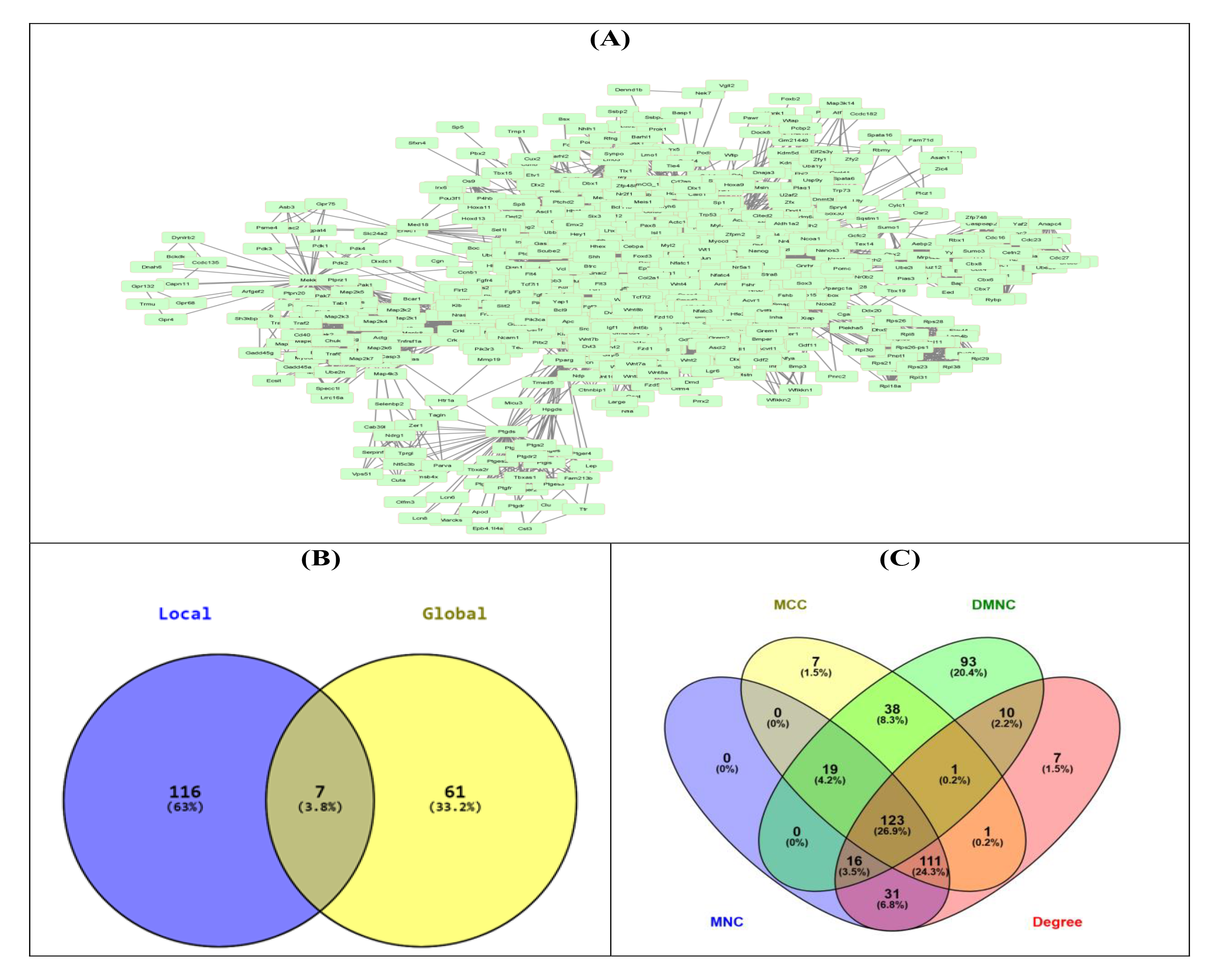
**[A]** protein-protein interaction network. **[B]**Venn diagram analysis for different topological analysis. **[C]** Venn diagram shows global and local analysis of network. Intersection of two circles represents overlapping genes representing both hub and bottleneck genes.

**Figure 3.**
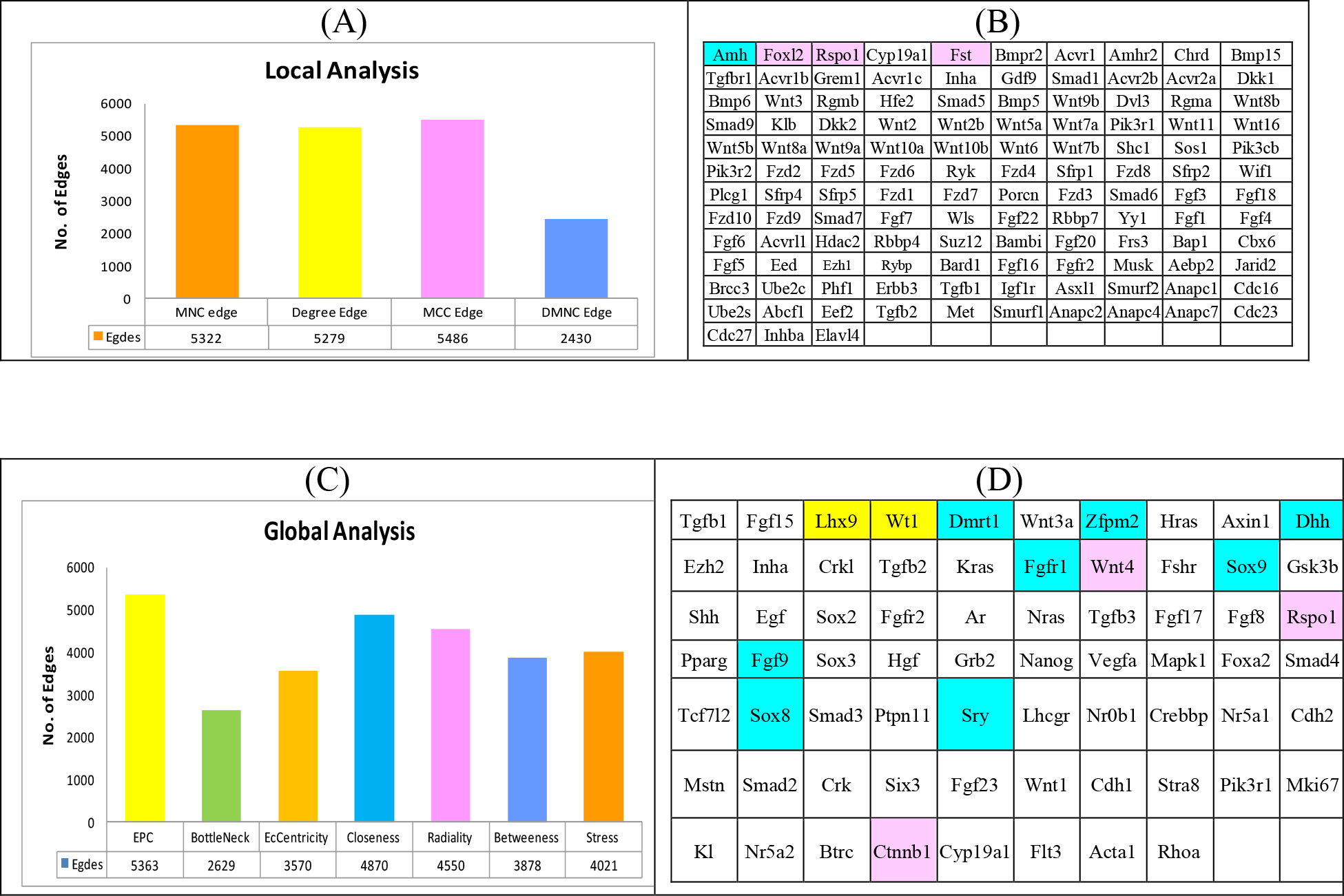
Cytohubba based topological Analysis **[A]** Based on local parameter **[B]** 123 genes common in both local based analysis and our total RNA RNAseq data **[C]** Based on global parameters **[D]** 68 genes common in both local based analysis and our total RNA RNAseq data

**Table 2.**
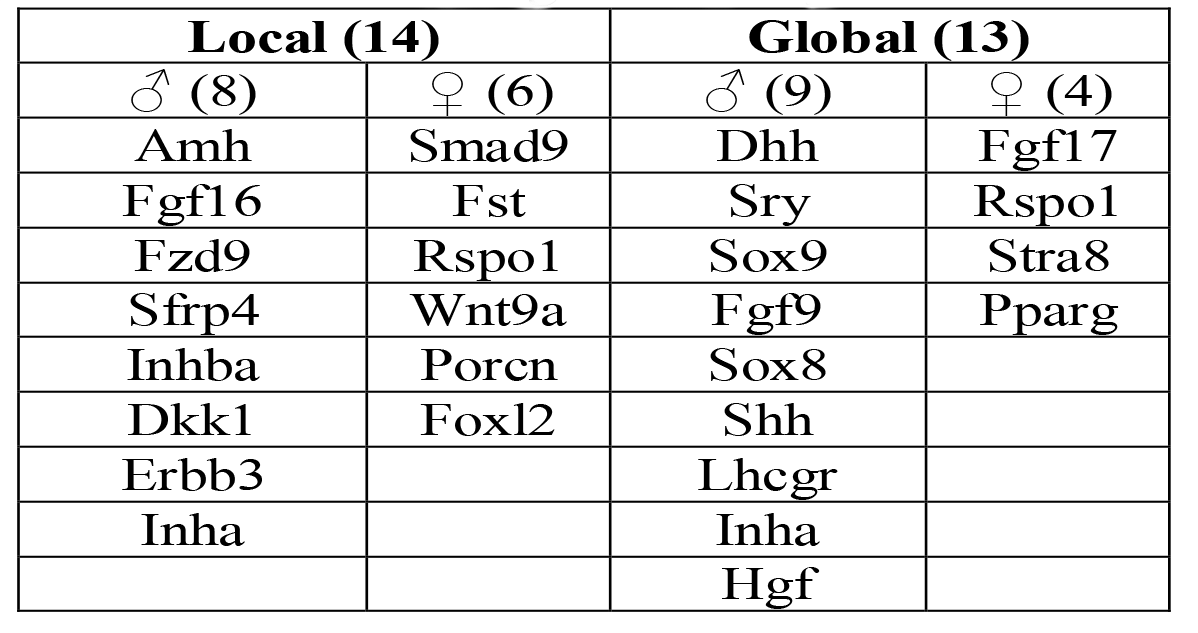
Differentially expressed genes from local and global analysis

### 3.3 Clustering and module analysis

The Cytoscape plugin MCODE v2.0.0 was applied to interpret closely interlinked regions from PPI network inform of clusters (modules) which was lacking through topological analysis. It predicts protein complexes in networks and represents relatively stable, multi-protein complexes that function as a single entity spatio-temporarily. Cluster formation is based on an algorithm that uses three-step processes: a high score is assigned to nodes whose neighbors are highly interacting; seed node i.e., high score nodes, recursively move out, adding nodes to complex that are above given threshold and post-processing applies filters to improve cluster quality (haircut and fluff). Thirty-seven clusters were obtained using MCODE with highly interconnected nodes and edges. The highest Mcode score obtained was 43.467 with 46 nodes and 1068 edges. For selection of important clusters out of 37, we arranged each cluster’s genes and overlapped them with our NGS data obtained earlier. Common genes with differential expression in male and female at any stage (11.5-13.5 dpc) were marked. Out of 37 clusters, 15 clusters were selected with differential gene expression common in data. Cluster 1 subnetwork with highest Mcode score, contained 5 DE genes. Cluster 2 subnetwork with 30.80 score possesses highest number of nodes and edges i.e., 133 and 2398 respectively. It had 9 DE genes in cluster. Cluster 3 did not have any DE genes therefore no considered for analysis. Next, cluster 4 had 2 DE genes in 13 nodes and 69 edges with 11.5 Mcode Score. Likewise, other clusters were also analysed with these parameters. From15 clusters, differential genes are further sorted into male and female-specific genes. Table 3 represents details of MCODE clusters with their scores, number of nodes, and edges, along with corresponding differentially expressed genes only.

**Table 3.** List of Clusters from PPIN and respective DEGs identified in male and female embryonic gonad using MCODE.

| Cluster No. | Nodes | Edges | MCODE Score | No. of Genes | Common Genes with Differential Expression Data (65) |  |
| --- | --- | --- | --- | --- | --- | --- |
|  |  |  |  |  | Male (49) | Female(16) |
| 1 | 46 | 1068 | 43.467 | 46 | Fzd9, Dkk1, Sfrp4 | Porcn, Wnt9a |
| 2 | 133 | 2398 | 30.803 | 133 | **Fgf9**, Fgf16, Cbx7, Fgf10 | **Bmp2**, Fgf17, **Fst**, Fgf2, Smad9 |
| 4 | 13 | 69 | 11.5 | 13 | **Ptgds** | Ptger3 |
| 5 | 31 | 193 | 9.7 | 31 | Hgf, Erbb4, Erbb3, **Amh** |  |
| 6 | 48 | 207 | 7.8 | 48 | Ptch1, Hhip, Lhcgr, Lhx1, Etv1, Nanos2 | Rnf43, Lgr6, Sp8 |
| 7 | 7 | 20 | 6.667 | 7 | Pak3, Pak7 |  |
| 8 | 29 | 93 | 6.286 | 29 | Uba1y, Gli1, Eif2s3y, Ddx3y, Kdm5d, Zfy2, **Sry**, Hsd3b1, Zfy1, Hsd17b3, Uty, Usp9y |  |
| 9 | 6 | 15 | 6 | 6 | Nphs2 |  |
| 10 | 36 | 124 | 5.657 | 36 | Shh, **Inha**, Inhba, **Sox9** |  |
| 11 | 16 | 110 | 5.6 | 16 | Insl3, **Sox8**, Cyp17a1, Cyp11a1, Star | Pparg, **Foxl2** |
| 13 | 32 | 107 | 5.548 | 32 | Actc1, Col2a1, Jag1 |  |
| 14 | 7 | 19 | 5 | 7 |  | Sohlh1, Figla |
| 16 | 6 | 11 | 4.4 | 6 | Pou3f2 | Pax7 |
| 17 | 7 | 13 | 6.33 | 7 | Ppargc1a |  |
| 24 | 3 | 3 | 3 | 3 | Vil1, Lgr5 |  |

### 3.4 Gene ontology and functional enrichment analysis

To characterize molecular mechanism of hub and bottleneck nodes involved in sex determination, we utilized DAVID online tool for functional enrichment analysis. To examine relative function of nodes we analysed different sets of nodes obtained from PPI network analysis. From topological analysis we examined nodes which were common in Global (68) and Local (123) analysis. Further, we separately analysed DE nodes from these Global (13) and Local (14) groups for differentially expressed genes. In addition to this we also performed functional enrichment analysis for nodes obtained from modular analysis which yielded DE genes present in different modules.

Ontology analysis of nodes obtained from Global analysis revealed a significant enrichment of gonadogenesis-related genes in different biological processes, including animal organ development with highest number of genes (65/68)tissue development (53/68 genes included), reproductive structure development, and reproductive system development with 35/68 genes. Sex determination term is also enriched for 30/68 genes, male sex differentiation with 24/68 genes. 60 genes out of 68 are involved in signaling processes which signifies importance of genes in global analysis and their involvement in crucial biological processes. Cellular component is also showing important terms like transcription factor complex, Wnt signalosome, cytoplasm, nucleus, RNA polymerase II transcription factor complex, etc. In molecular function, terms are enriched like receptor binding, transcription factor binding, growth factor activity, beta-catenin binding, chromatin binding, FGF receptor binding, etc. the differentially expressed global nodes (13) were enriched with 600 BP ontology terms including male sex differentiation, reproductive structure development, reproductive processes, etc. CC is also enriched with terms like extracellular space, transcriptional factor complex, etc., and MF like receptor binding, growth factor activity, protein binding, FGF receptor binding, and many more. In KEGG pathway and wiki pathway analysis, terms enriched were mostly involved in pathways related to sex determination while a few terms were related to cancer pathways, which makes these genes very crucial in not only gonadogenesis but also in related disorders.

The ontology analysis of Local parameters-based nodes (123) revealed that these nodes are involved in BP like, cell surface receptor signaling pathway, cellular response to growth factor stimulus, canonical Wnt signaling pathways, animal organ morphogenesis, and regulation of developmental processes.CC terms linked to genes are PcG protein complex, transferase complex, extracellular space, nuclear ubiquitin ligase complex, and many more. The molecular function included frizzled binding, receptor binding, Wnt-protein binding, growth factor activity, etc. Terms like signal transduction, WNT ligand biogenesis, trafficking, downstream signaling of activated FGFR2 and many others were enriched in Reactome pathways. Differentially expressed nodes of Local analysis (14), were expressed in BP terms like cell surface receptor signaling pathway, canonical Wnt signaling pathway, female, gonad development, and development of primary female sexual characteristics indicating an important involvement of these genes in ovary maintenance.CC and MF also indicate direction of female gonad development as terms enriched are extracellular space, inhibin A complex, etc., in CC and receptor binding, Wnt-protein binding, inhibin binding, receptor antagonist activity, and many others. Along with involvement in gonadogenesis pathways terms, KEGG pathways are also showing terms related to breast cancer, gastric cancer, and other diseases.

We further conducted functional enrichment analysis for differentially expressed nodes obtained from modular analysis. Out of 37, 15 modules had differentially expressed genes. Therefore, to analyze importance of clusters we conducted gene ontology analysis of top three clusters with highest scores and differential genes. Results show that cluster 1 is mainly associated with biological processes like Wnt signaling pathways, cell-cell signaling and animal organ development. Cluster 2 genes are mainly enriched with terms like protein metabolic processes, response to growth factors and gene expression. Cluster 4 is mainly involved in prostaglandin metabolic process, fatty acid biosynthetic processes.

From15 modules, 49 nodes were male-specific and 16 nodes were female-specific. For biological processes, male-specific nodes were enriched in terms like male-sex differentiation, animal organ development, and other terms related to gonadogenesis. CC includes cell body, nucleus, intracellular part etc., terms while MF includes growth factor activity, receptor binding, FGF receptor binding, organic cyclic compound binding, etc. Ovarian steroidogenesis, basal carcinoma, and pathways in cancer are a few top terms enriched in KEGG pathways.

The biological processes of female-biased nodes show terms like connective tissue development, anatomical structure development, positive regulation of gene expression, developmental processes, etc.CC shows terms like transcription factor complex, extracellular space, nucleus, etc. MF enriches terms like, RNA polymerase II core promoter proximal region sequence-specific DNA binding, core promoter proximal region sequence-specific DNA binding, receptor binding, and many more. KEGG pathway terms are related to cancer pathways, Wnt signaling pathways, TGF-beta signaling pathways, and calcium cell signaling pathways.

### 3.5 Gene-miRNA network analysis

We aimed to predict a few differentially expressed miRNAs (DEmiRs) depicted in network that could potentially regulate key genes during gonadogenesis. To analyze interaction between miRNA and their targets we created an interaction network of 25 selected genes and miRNAs targeting them. Using miRDB database for each gene, a list of miRNAs was obtained. We considered only those miRNAs whose target scores were above or equal to 80. Out of 25 selected genes database could find miRNAs targeting only 23 genes while for two genes Sry and Ptgds no miRNAs were enlisted. To visualize interaction network, genes and their miRNAs were exported to Cytoscape (Figure 6.4) where each gene was connected to their unique miRNAs and a few common nodes and edges were interconnected forming a complex network.

Further, list of total miRNAs obtained from miRDB overlapped with our small RNAseq data. We obtained miRNAs that are differentially expressed in male (26 miRNAs) and female (16 miRNAs) embryonic gonads. These miRNAs were taken to miRTarBase and their targets were extracted. These targets were again overlapped with our transcriptomic data which revealed targets that were differentially expressed in males and females. miRNAs that are upregulated in males, target genes that are downregulated in females and vice versa might be important in this interlaced connection that governs gonadogenesis (Figure 5).

**Figure 4.**
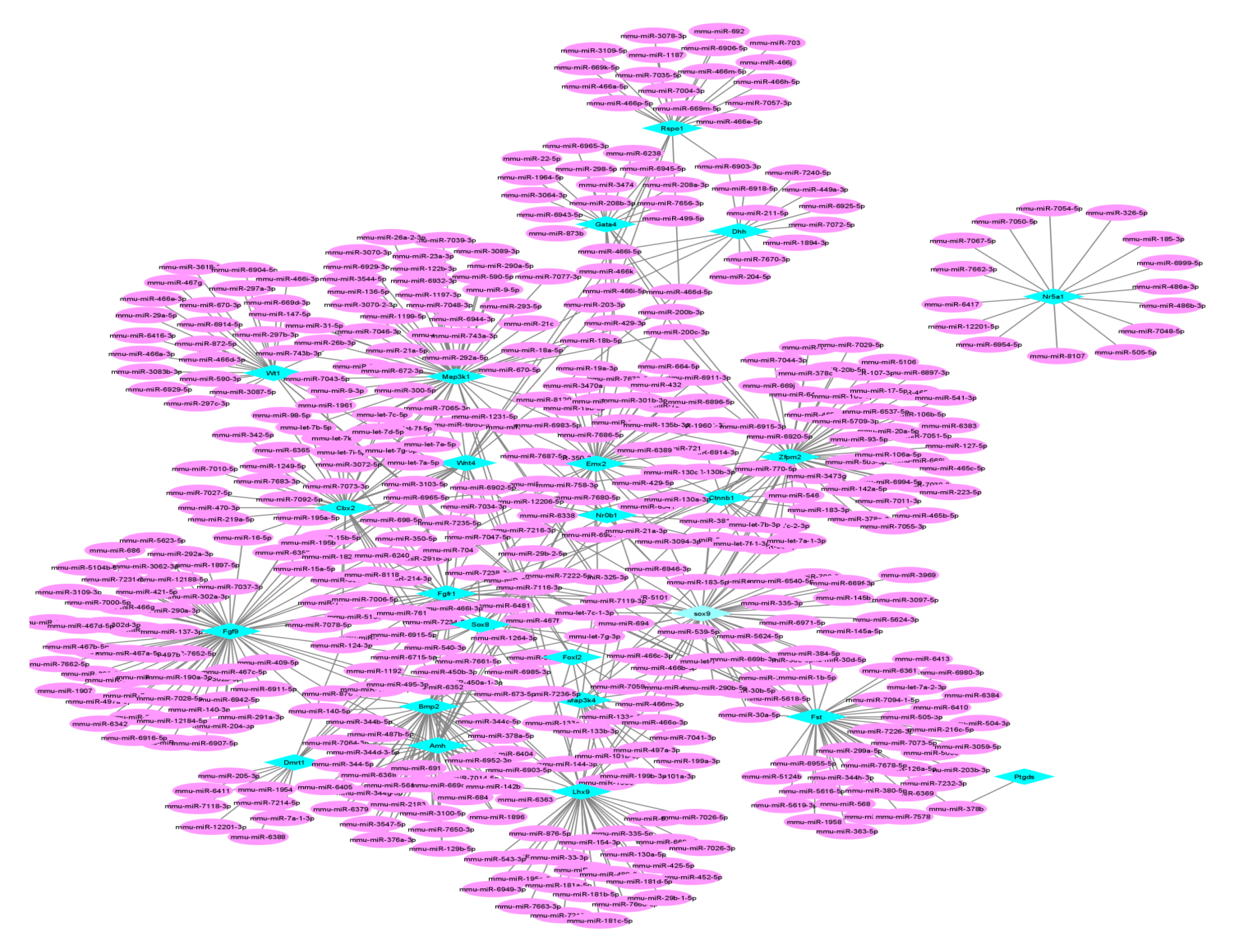
Gene-miRNA interaction network. The interaction was visualized on Cytoscape. Sphere corresponds tonumber of interactions they are involved. Genes under query are in sky blue and miRNAs targeting them are pink in colour.

**Figure 5.**
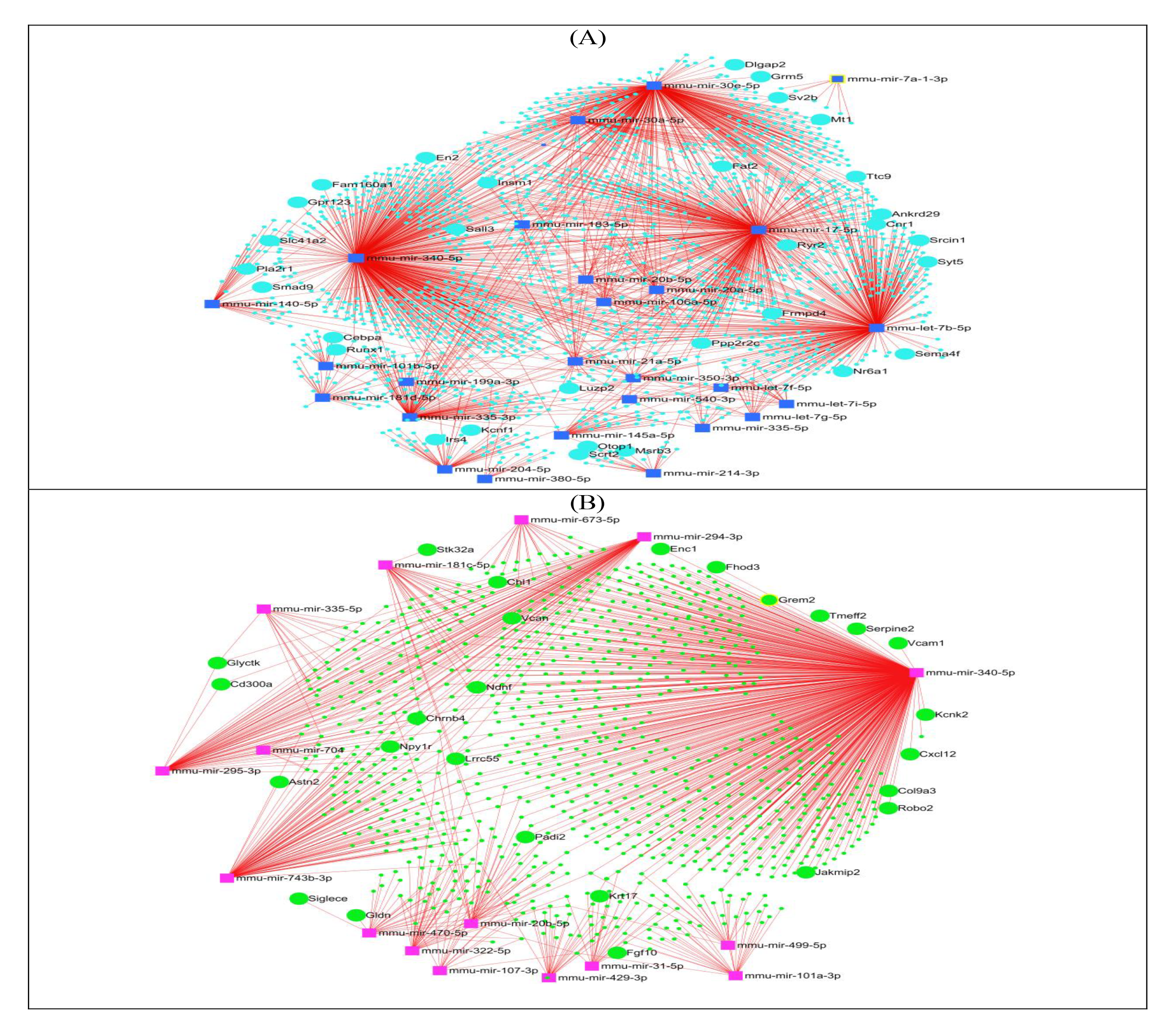
Male **[A]** and Female **[B]** specific network of differentially expressed miRNAs from main network and their targets with relativelly lower expression in corresponding sex type. Navy blue and pink squares depicts male and female differentially expressed miRNAs while light blue and green circles shows differentially expressed targets. Very small blue and green dots represents targets which are not differntially expressed.

To understand functional involvement of differentially expressed targets of differential miRNAs we performed functional enrichment analysis using DAVID. Male-specific miRNAs targeting female-specific targets (31) Figure 6 a. were enriched in biological process terms like regulation of localization, embryonic organ development, and negative regulation of biological processes, developmental processes, and cell differentiation. A few CC in which these nodes are enriched is cell projections, synapses, and transcription factor complexes. MF terms are mainly related to DNA binding and transcription factor activities. On the other hand, targets of female-specific miRNAs are mainly enriched in biological processes like cell adhesion, system development, animal organ development, developmental processes, and biological regulation. Cellular components revealed are cell surface, cell periphery, plasma membrane, and proteinaceous extracellular matrix. KEGG pathways include glycosaminoglycan binding, carbohydrate derivative binding, and binding, etc.

**Figure 6.**
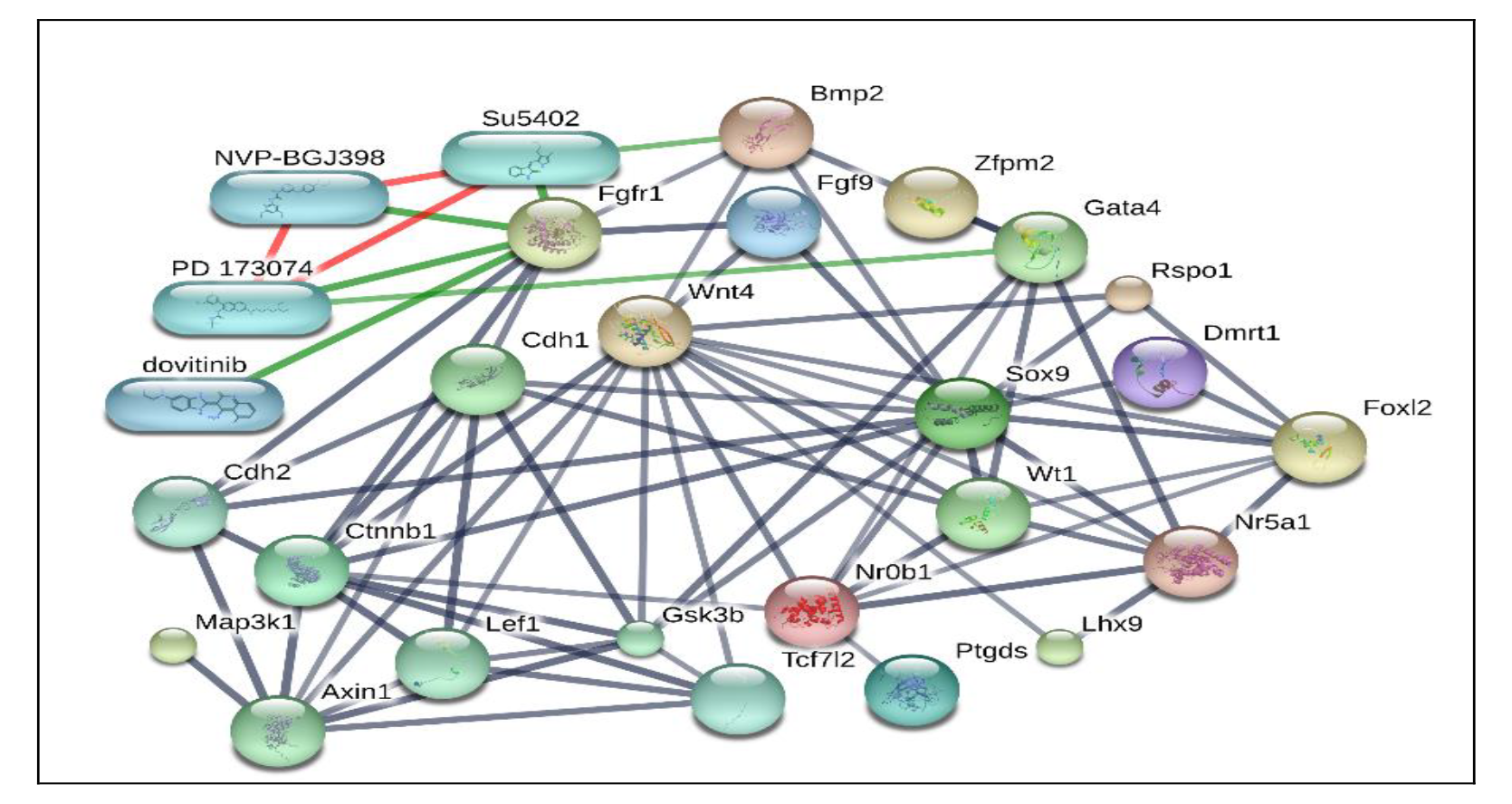
Chemical-Protein interactions graph generated using STITCH tool. STITCH tool was queried using 25 selected genes. Nodes, which were not directly related were removed in order to highlight the most relevant concepts to repositioned drug.

### 3.6 Prediction of lncRNA and their interaction with sex-determining genes

This study was performed to identify novel lncRNA that could modulate molecular mechanism of sex determination. LncRNA has a wide range of regulatory roles in gene expression through up regulation or down-regulation of target genes. A comprehensive lncRNA-mRNA interaction prediction is necessary for estimation and validation of lncRNA function that can regulate biological processes. Previously insearch for these regulators Chen et al 2012, conducted a study to identify ovay-spcecific biomarkers. Microarray assay yielded 56 nc-RNAs showing higher expression from 11.5 −13.5 dpc in embryonic mouse ovary. Out of 11 validated candidates displaying sexually dimorphic expression, they found oncRNA3 lncRNA, an antisense tocoding gene Pank1, predominantly expressed in ovarian somtic cells. Keeping into consideration importance of lncRNA in epigenetic modification we analysed their interaction with selected genes using a robust LncRNA prediction tool. Based oninteraction energy lncRRIsearch online tool predicts all possible query-target pairs based on two criteria: minimum interaction energy and a sum of all interaction energies of local base-pairing forRNA-RNA pair. Tool provides a list of top 100 lncRNA of lowest sum of energy, below threshold. From list, we picked only differentially expressed lncRNA in embryonic gonads from 11.5-13.5 dpc using Jameson Database (Jameson et al, 2012). Table shows alldifferentially expressed lncRNA for specific targets below threshold energy which was set to −16 kcal/mol.

**Table 4.** Male and female specific LncRNA targeting key genes of sex determination and gonad differentiation

| Target gene | lncRNA (Transcript name) | Sum of energy | Minimum energy | Number of local base-pairing interactions | Expression of lncRNA transcript (Jameson database, 2012) |
| --- | --- | --- | --- | --- | --- |
| **Bipotential genes** |  |  |  |  |  |
| Cbx2 | ENSMUST00000136990(Emx2os-201) | -414.04 kcal/mol | -23.78 kcal/mol | 22 | Enriched in female supporting cells at 11.5 and 13.5 dpc. |
| Emx2 | ENSMUST00000136990(Emx2os-201) | -1329.05 kcal/mol | -214.76 kcal/mol | 35 | Enriched in female supporting cells at 11.5 and 13.5 dpc. |
| **Testis-specific** |  |  |  |  |  |
| Sry | ENSMUST00000150671(2010204 K13Rik-203) | -861.83 kcal/mol | -30.78 kcal/mol | 40 | Enriched in female supporting cells at 12.5 dpc. |
|  | ENSMUST00000155906(Gimap1os-201) | -279.86 kcal/mol | -25.83 kcal/mol | 14 | Enriched 12.5 endothelial cells enriched female |
| Sox9 | ENSMUST00000136990(Emx2os-201) | -1603.04 kcal/mol | -35.70 kcal/mol | 68 | Enriched in female supporting cells at 11.5 and 13.5 dpc. |
| Sox8 | ENSMUST00000136990(Emx2os-201) | -332.84 kcal/mol | -18.88 kcal/mol | 19 | Enriched in female supporting cells at 11.5 and 13.5 dpc. |
| **Ovary-specific** |  |  |  |  |  |
| Wnt4 | ENSMUST00000125191(Tctn2-202) | -1056.07 kcal/mol | -81.98 kcal/mol | 24 | Enriched in male supporting cell from 12.5-13.5dpc |
|  | ENSMUST00000206519(Snhg1-215) | -592.75 kcal/mol | -58.20 kcal/mol | 26 | At 11.5 dpc it is enriched in germ cells in both sexes. By 12.5 dpc increases in male germ cells and downregulates in male supporting cell. While by 13.5 it is left only in male germ cells. |
| Rspo1 | ENSMUST00000136990(Emx2os-201) | -510.05 kcal/mol | -31.75 kcal/mol | 20 | Enriched in female supporting cells at 11.5 and 13.5 dpc. |

### 3.7 Protein-Drug interaction network

In order to link drugs associated with input proteins, we utilized STITCH a chemical-protein interactions database. Network consists of 27 nodes and 67 edges. Network had significantly more interactions (67 edges) than expected (9 edges) Figure 6. This is mostly because our protein has more interactions among themselves with respect to other proteins of similar size from genome. This indicates that proteins are to some extent biologically connected, to form a group. STITCH has added 10 partners to show networks around 25 input genes. It included six genes Cdh1, Cdh2, Axin1, Gsk3b, Lef1, Tcf7l2, and four drugs Su5402, PD173074, NVP-BG-J398 and dovitinib. Interestingly, both Cdh1/2 (Okamura et al. 2003; Piprek et al., 2019), Gsk3b, and Lef1 (Garcia-Moreno et al., 2019) are related to sex determination. Out of 25 genes twenty-one genes were found to interact with other participants’ of network. However, only three genes were directly interacting with drugs present in network. FGFR1 is forming common connections with all drugs while along with FGFR1, BMP2 is interacting with Su5402 and GATA4 is interacting with PD173074.

### 3.8 Disease association and Dragome Analysis

Literature survey was conducted to search relation between selected genes and disease and disorders related to sex determination and reproductive organs. Table 5 shows congenital disorders related to genes of interest. Table also highlights diseases related togonadal diseases in adults. To analyse disease-drug association it is necessary to analyses duggable properties of a protein. Predicting Druggability i.e., ability of a protein to bind drug-like molecules with high affinity of a protein is of utmost importance during target identification during drug discovery phase. DrugnomeAI was utilized to predict druggability of targets for crucial drug target selection. From25 selected genes eleven genes fall under criteria of low probability while fourteen were showing very high probable chances of druggability. The fourteen includes Bmp2, Wt1, Fgf9, Soz9, Ptgds, Amh, Fgfr1, Dhh, Ctnnb1, and Wnt4, rspo1, Fst, Map3k1 and map3k4. Further to recheck this data we also utilized PockDrug-server. It returns an average druggability probability of different pockets ranking it as a potentially or difficult druggable pocket. Out of 25 selected candidates, only 12 were found to have a probability of use as a potential target for drug designing purpose. These 12 candidates are: Wt1, Nr5a1, Gata4, Fgf9, Fgfr1, Sox9, Sox8, Ptgds, Ctnnb1, Wnt4, Fst, Bmp2, and Amh.

FGF9 (1IHK) and PTGDS (4ORUA) are selected for further analysis as only these fulfil two criteria i.e., probably good druggability score and full-length structure. Pockets are estimated as displayed in Figure 7. In 1IHK one pocket (P0) with druggability probability of 0.86 (±0.03) and two pockets (P1 and P2) with druggability probability of 0.08 (±0.01) and 0.51 (±0.01) respectively which have small pockets with residues less than 14 were predicted. Similarly, in 4ORUA also one pocket (P0) with a druggability probability of 0.87 (±0.02) and two pockets (P1 and P2) with a druggability probability of 1.0 (±0.0) and 0.49 (±0.15) respectively with small pockets (residues less than 14) were predicted. For both proteins, P0 are favorable pockets as druggables and can be utilized for further analysis.

**Figure 7.**
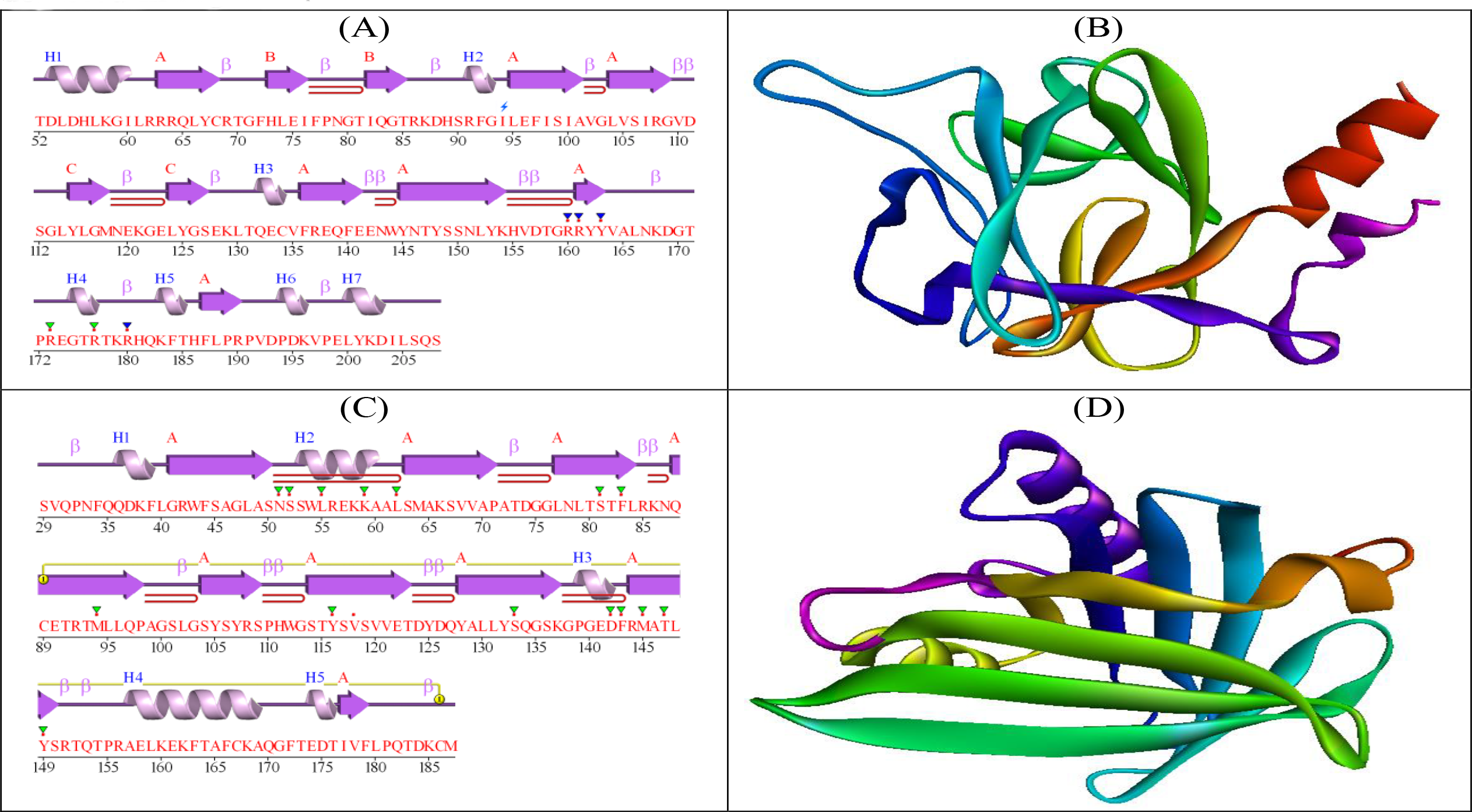
**2D and 3D structure visualization [A]** Secondary structure of FGF9 showing helices labelled as H1-H7, Beta sheets, and beta turns. **[B]** 3D structure of FGF9 **[C]** Secondary structure of PTGDS showing helices, Beta sheets, and beta turns. **[D]** 3D structure of PTGDS

**Table 5.**
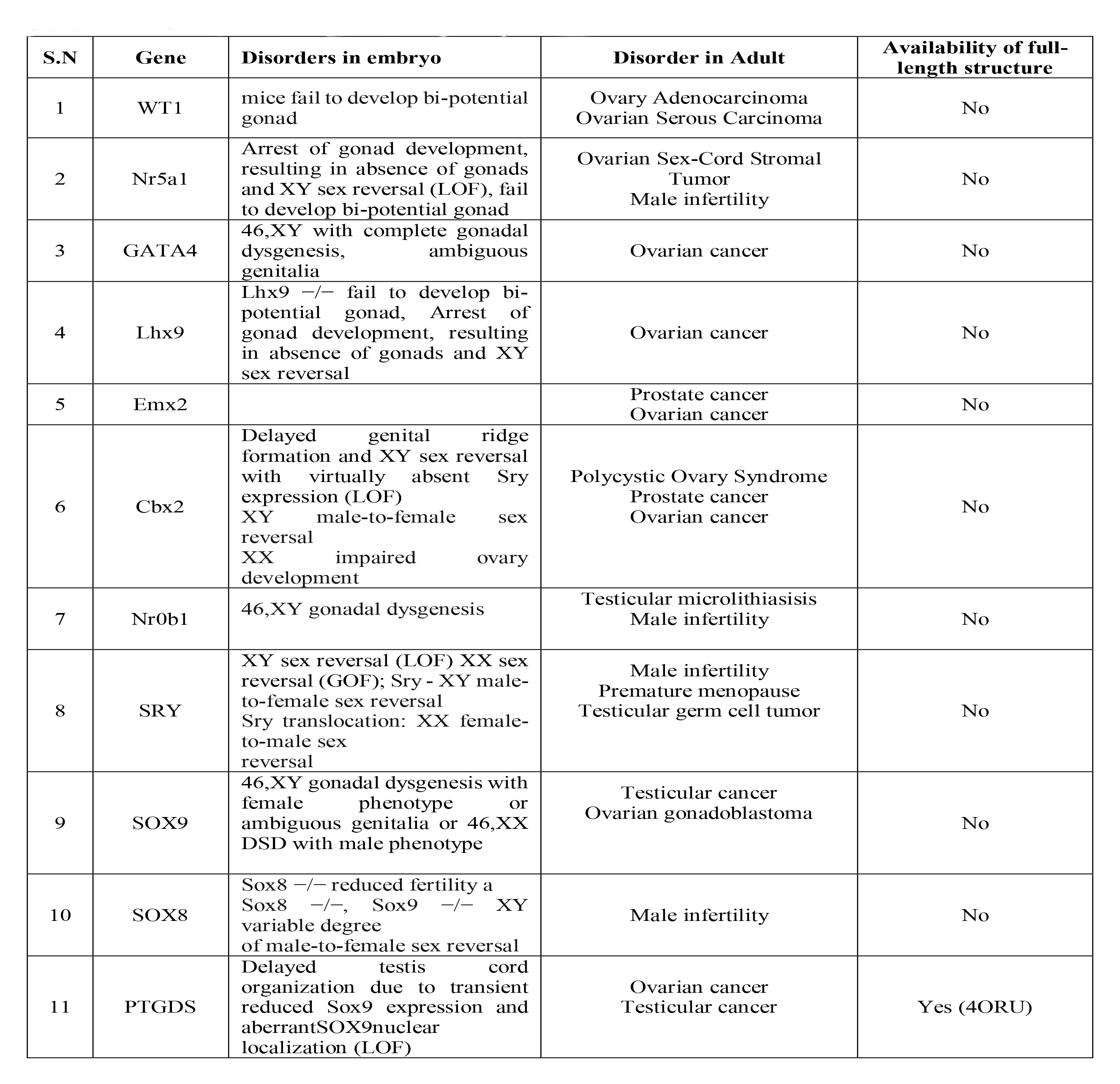
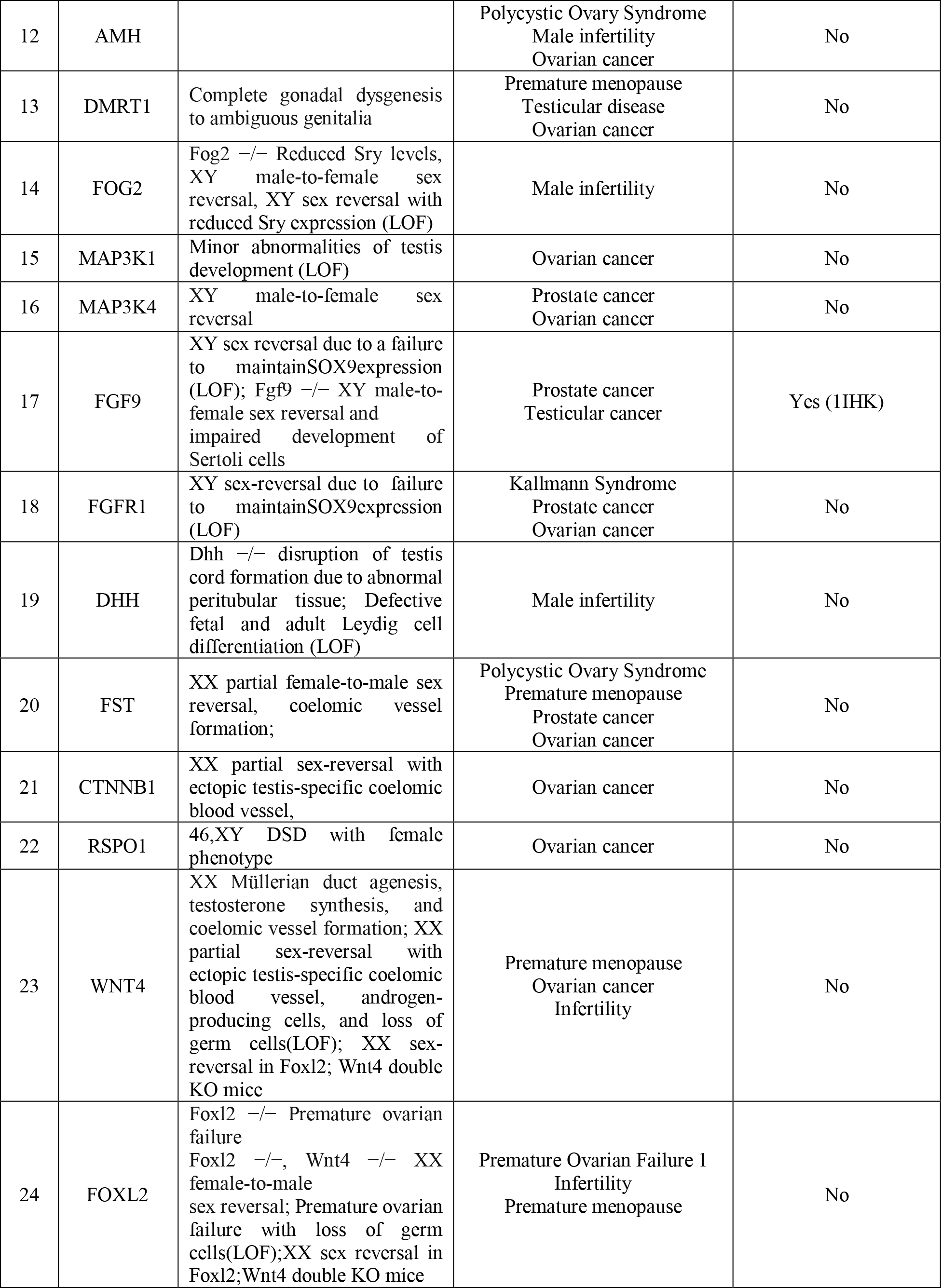
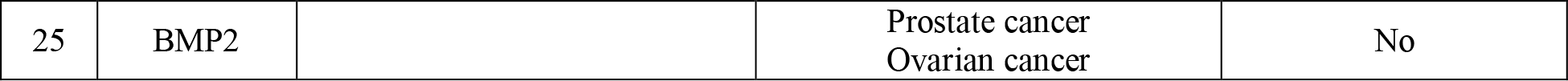
Gene-disease association table for targeted protein

### 3.9 Receptor Selection

Out of twelve proteins analyzed through Pockdrug tool only two candidates, FGF9 and PTGDS, had their full-length structure available on UniProt and were found to be suitable for further drug designing. The receptor FGF9 (PDB ID: 1IHK) and PTGDs (PDB ID: 4oru) were selected and retrieved for further analysis.FGF9 sequence consists of 208 amino acids (https://www.uniprot.org/uniprot/P31371). Structure selection was performed by X-ray crystallographic method at a resolution of 2.20 Å. topological analysis of protein-protein interaction network established in this study, predicts Fgf9 and ptgds as both hub and bottleneck. FGF9 and PTGDs are involved in various biological activities and are one of most essential elements in sex determination and testicular development. Its involvement in many disorders and diseases with least known treatments made these proteins perfect candidates for this study. To identify new compounds that could potentially inhibit FGF9 and PTGDS through binding to active sites of protein, virtual screening and processing are carried out through various bioinformatics tools. Out of all crystal structures of FGF9 and PTGDS in PDB one without any complex or dimer, with least resolution was selected. Virtual screening and validation of receptor structure are depicted in Figure 7.

### 3.10 Structure Validation

Target receptor structures were prepared using various tools like Modrefiner and SAVES (VERIFY 3D, ERRAT and PROCHECK). ERRAT program is used for verifying protein structures determined by crystallography. Error-values are plotted as a function of position of a sliding 9-residue window. Figure 8 provides Errat validation and quality check for (A) FGF9 and (B) PTGDS. Stereochemical quality and accuracy of 3D structure were evaluated after refinement process using the Ramachandran Map calculation with PROCHECK program. Details of Ramachandran plot statistics of FGF9 are shown in Figure 9. (A). Indeed, in Ramachandran plot, residues were classified according to their regions in quadrangle. Red regions in graph indicate the most allowed regions, whereas yellow regions represent allowed regions. Glycine is represented by triangles and other residues are represented by squares. Ramachandran plot phi and psi angles contributing to conformation of amino acids excluding glycine and proline with 91.9% (124 amino acids) residue in most favored region, 8.1% (11 amino acids) in additional allowed region, 0% (0 amino acids) in generously allowed region and disallowed region. Number of glycine and proline residues was 14 and 6 respectively. Similarly, Figure 9 (B) shows PTGDS Ramachandran Plot with 92.1% (128 amino acids) residue in most favored region, 7.9% (11 amino acids) in additional allowed region, 0% (0 amino acids) in generously allowed region and disallowed region. These plots indicate that structure is of best quality.

**Figure 8.**
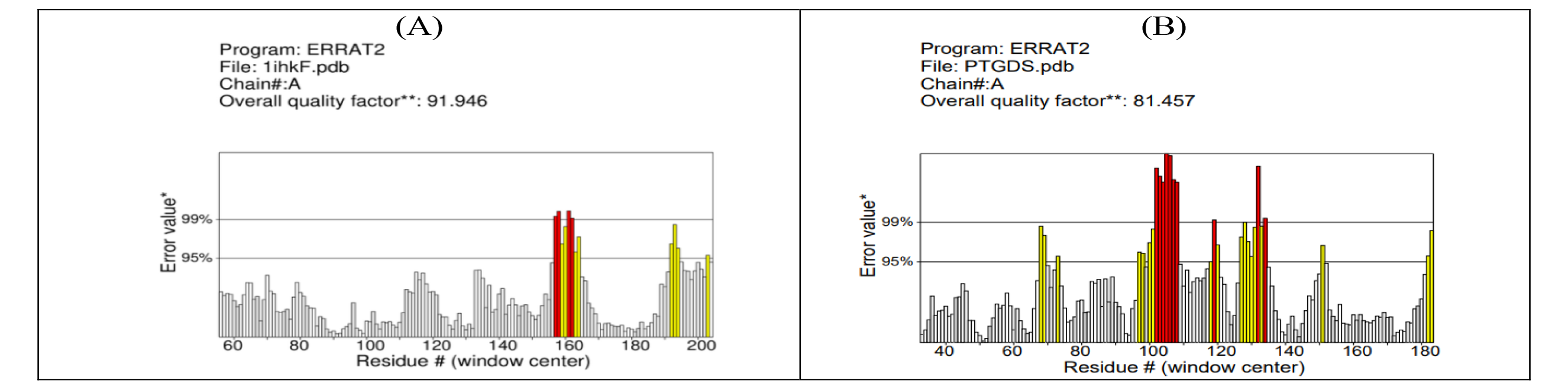
ERRAT2 validation: verification of protein structure and quality check. [A] FGF9 [B] PTGDS

**Figure 9.**
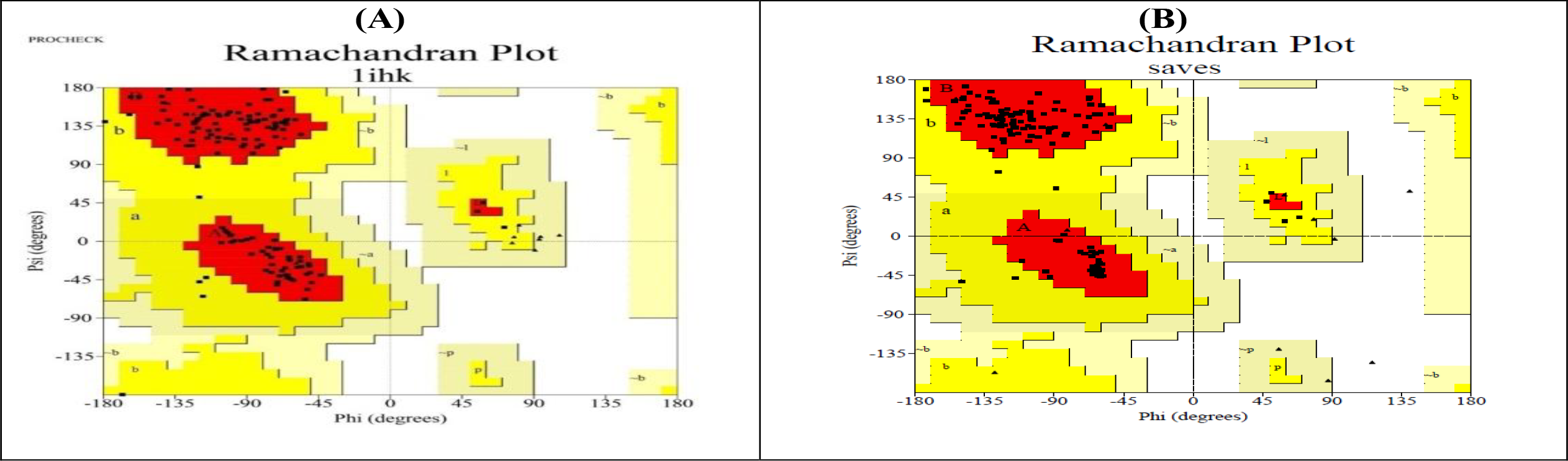
**Receptor structure validation** Ramachandran plot of receptor **[A]** Fgf9 & **[B]** Ptgds generated from Procheck server.

### 3.11 Active Site Analysis

Active site and their prominent residues were identified using BIOVIA Discovery Studio 2019 and POCASA respectively. Common active site region obtained from both tools/servers was taken for analysis, which was showing best binding site further used for docking calculation using LibDock and CDOCKER. POCASA (POcket-CAvity Search Application) implements a geometric search algorithm called Roll, which uses both grid system and probe sphere (sphere-based grid) to search pockets and cavities. Region between protein and probe surface and those surrounded by protein surface are called pockets and cavities, respectively. Further, noise points were removed and pockets were ranked. Figure 10. (A) and (B) represents common active site of FGF9 and PTGDS. Dots represent active site residues identified from POCASA showing five active sites while sphere represents sites identified by Biovia. Common sites are marked in yellow dots as shown inFigure 10 for both protein structures.

**Figure 10.**
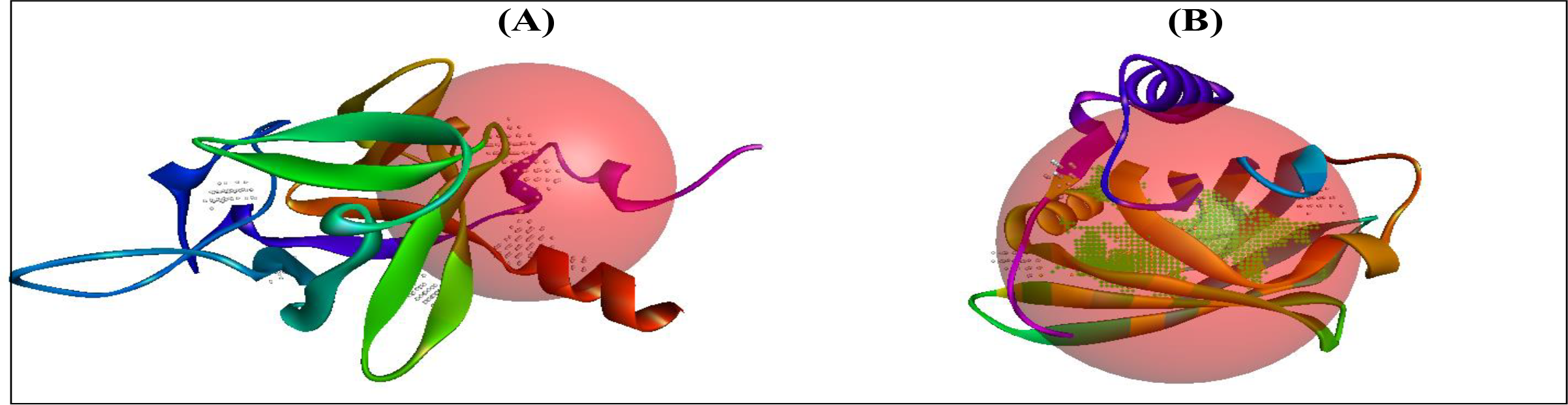
**(A)** Active site identification using BIOVIA Discovery Studio 2019 in sphere green + symbols + Ligand-binding pocket identification using POCASA in Pocket A in yellow dotted others pockets B, C, D, and E showing grey in color. **(B)** Active site residue identification of POCASA pocket A result was calculated by BIOVIA Discovery Studio 2019.

### 3.12 Ligand Selection

Docking predicts experimental binding affinities of small molecules and ligands within binding site of particular receptor targets. Identified target protein FGF9 play important role in various biological processes including follicular development, ovarian follicular cysts, and testicular tumorigenesis. On other hand, PTGDS is also involved in cellular invasion in testicular cancer cells, ovarian cancer, Prostate cancer, and Adult Syndrome (). For retrieval of new compounds that could potentially inhibit FGF9 and PTGDS from interacting with other molecules in pathways through active binding pockets, virtual screening was carried out using LibDock a module of Discovery Studio. These drugs were retrieved from Drugbank database and are FDA approved. In total 9213 drugs were retrieved from Drugbank of which 7407 drugs were further analysed whose molecular weight was between 200-600 gm/mol. After removal of all conformers, we are left with 3736 compounds for FGF9 and 2004 for PTGDs. After Libdock and CDocker analysis, 236 (FGF9) and 181 (PTGDS) drugs were left. Further after ADME and TOKAT analysis, we were left with 63 and 38 drugs for both FGF9 and PTGDS respectively. Finally, from these 16 and 14 drugs were shortlisted for MD simulations and MMGBSA (Figure 11), (Table 6 and 7).

**Figure 11.**
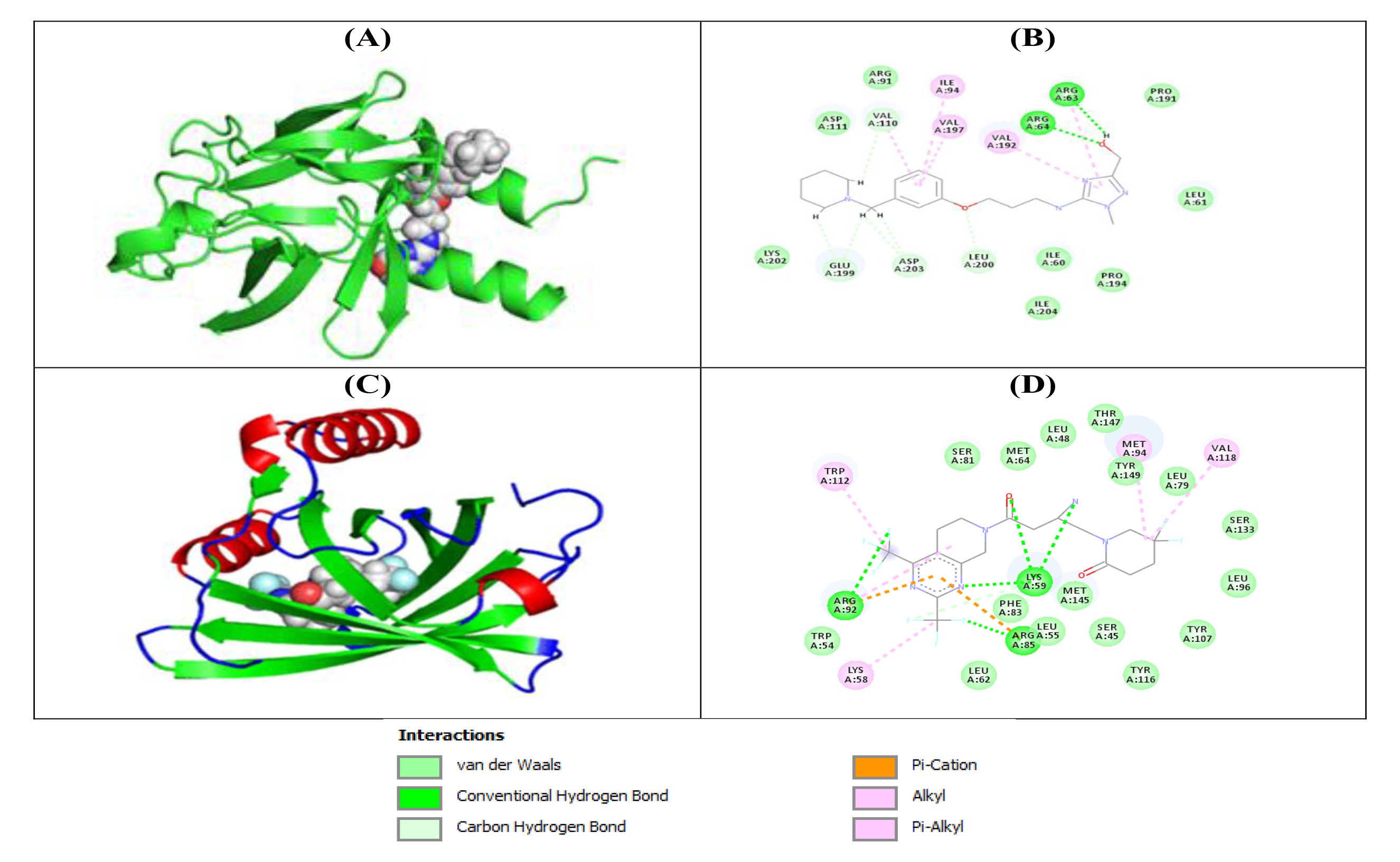
**A:** 2D representation of CID: interactions with FGF9 **B:** molecular interaction of DB12884 and H-Bonds details with FGF9. **C** 2D representation of interactions with Ptgds **B:** molecular interaction of DB12412 and H-Bonds details with PTGDS.

**Table 6.** Top ligands docking with FGF9. Table showing docking scores, ADME properties and toxicity prediction

| CID Number | DB ID | Drug group | Molecular weight | Libdock Score | Cdocker Interaction | ESOL Class | GI absorption | BBB permeant | Pgp substrate | CYP2D6 inhibitor | Lipinski violations | TOPKAT |
| --- | --- | --- | --- | --- | --- | --- | --- | --- | --- | --- | --- | --- |
| 198757 | DB06506 | Investigational | 400.146 | 141.833 | 44.5741 | Moderately soluble | High | No | Yes | Yes | 0 | Non-mutagen |
| 9905633 | DB06106 | Investigational | 332.16 | 149.509 | 40.1042 | Very soluble | High | No | No | No | 0 | Non-mutagen |
| 55473 | DB12884 | Investigational | 359.232 | 155.073 | 42.4375 | Soluble | High | No | Yes | No | 0 | Non-mutagen |
| 9675990 | DB13048 | Investigational | 392.221 | 131.488 | 49.7606 | Moderately soluble | High | Yes | No | Yes | 0 | Non-mutagen |
| 24798097 | DB08018 | Experimental | 373.203 | 155.158 | 41.9136 | Soluble | High | No | Yes | Yes | 0 | Non-mutagen |
| 25021186 | DB08019 | Experimental | 373.203 | 157.94 | 41.1598 | Soluble | High | No | Yes | Yes | 0 | Non-mutagen |
| 5326866 | DB08064 | Experimental | 399.146 | 140.859 | 40.2365 | Moderately soluble | High | No | No | No | 0 | Non-mutagen |
| 448271 | DB07416 | Experimental | 302.209 | 135.721 | 41.2318 | Soluble | High | Yes | No | No | 0 | Non-mutagen |
| 55483 | DB12313 | Approved; investigational | 356.246 | 161.243 | 48.137 | Soluble | High | Yes | Yes | Yes | 0 | Non-mutagen |
| 30949 | DB13681 | Experimental | 316.226 | 138.515 | 47.438 | Moderately soluble | High | Yes | No | Yes | 0 | Non-mutagen |
| 28892 | DB13731 | Experimental | 307.215 | 137.956 | 44.0952 | Soluble | High | Yes | No | Yes | 0 | Non-mutagen |
| 5794 | DB09350 | Approved; vet. approved | 338.209 | 146.776 | 44.0062 | Moderately soluble | High | Yes | No | Yes | 0 | Non-mutagen |
| 4031 | DB12554 | Approved; investigational | 429.252 | 151.741 | 41.5944 | Moderately soluble | High | Yes | Yes | Yes | 0 | Non-mutagen |
| 6914657 | DB12731 | Investigational | 391.226 | 147.315 | 41.176 | Moderately soluble | High | Yes | No | Yes | 0 | Non-mutagen |
| 5105 | DB08806 | Experimental | 348.205 | 146.905 | 41.0265 | Soluble | High | Yes | Yes | Yes | 0 | Non-mutagen |
| 6420121 | DB08259 | Experimental | 270.194 | 127.609 | 43.1656 | Soluble | High | Yes | Yes | No | 0 | Non-mutagen |

**Table 7.** Top ligands docking with PTGDS. Table showing docking scores, ADME properties and toxicity prediction

| CID | DB ID | Molecular weight | Libdock Score | Cdocking Interaction | ESOL Class | GI absorption | BBB permeant | Pgp substrate | CYP2D6 inhibitor | Lipinski #violations | TOPKAT |
| --- | --- | --- | --- | --- | --- | --- | --- | --- | --- | --- | --- |
| 447371 | DB02008 | 463.504 | 120.215 | 53.9053 | Moderately soluble | High | No | Yes | No | 0 | NON mutagen |
| 447219 | DB07548 | 432.399 | 122.373 | 48.5815 | Moderately soluble | High | No | No | Yes | 0 | NON mutagen |
| 1.2E+07 | DB12412 | 489.37 | 122.26 | 50.2993 | Moderately soluble | High | No | No | Yes | 0 | NON mutagen |
| 65906 | DB12523 | 432.503 | 123.159 | 42.7598 | Moderately soluble | High | No | No | No | 0 | NON mutagen |

### 3.13 Docking Calculation

CDOCKER is a CHARMm-based docking tool (Wu et al. 2003). A set of ligand compound conformations were generated using high-temperature molecular dynamics, with different random seeds. LibDock protocol is an interface to LibDock program developed by Diller and Merz (Diller and Merz, 2001). LibDock uses protein site features referred to as HotSpots. Hot Spots consist of two types: polar and apolar. A polar Hotspot is preferred by a polar ligand atom (for example a hydrogen bond donor or acceptor) and an apolar HotSpot is preferred by an apolar atom (for example a carbon atom). Receptor HotSpot file is calculated prior to docking procedure. Figure 12 (A-B) is showing docking interaction of FGF9 with its complex and Figure 12 (C-D) interaction of PTGDS with its compounds.

**Figure 12.**
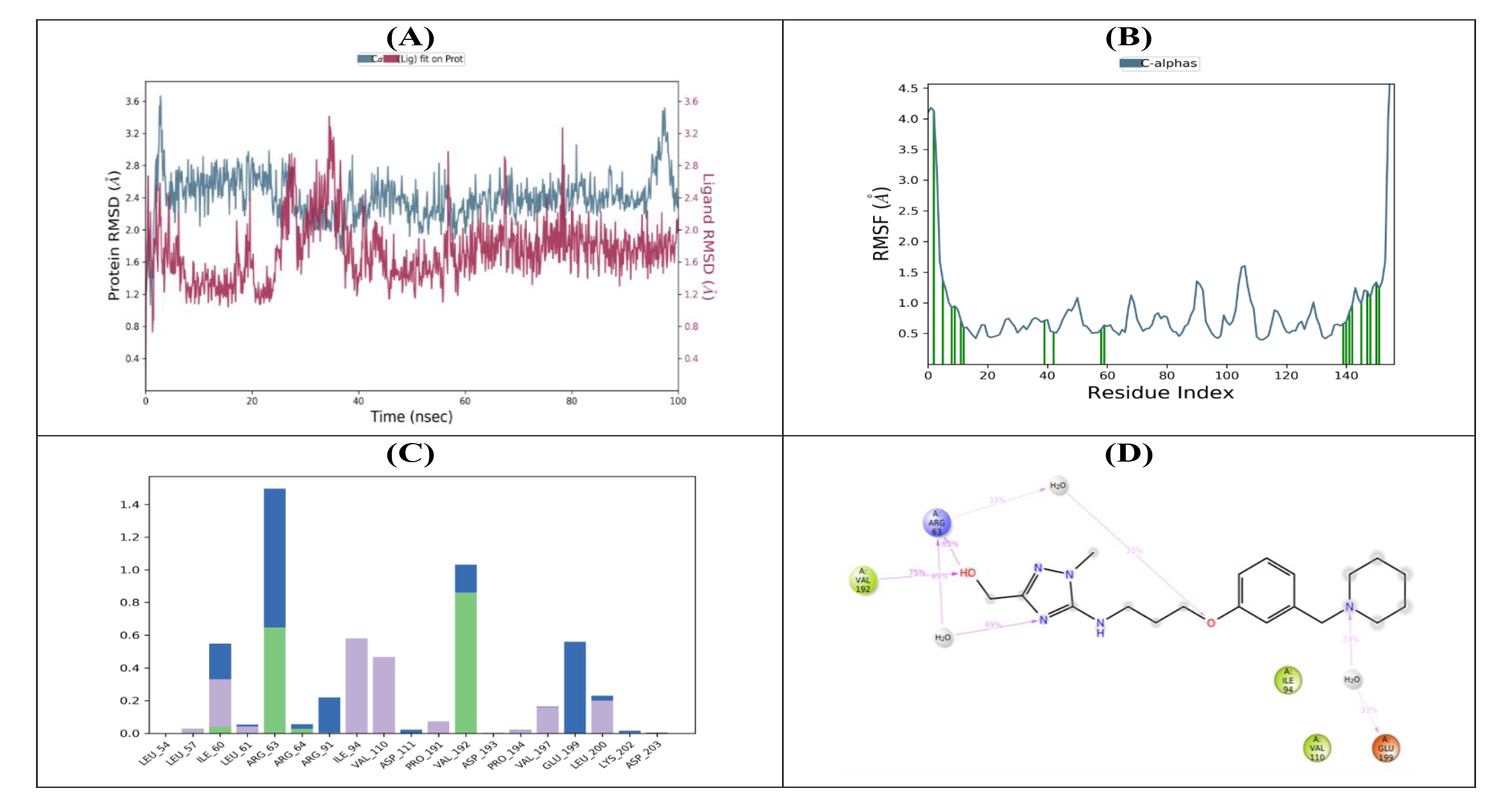
**Results of MD simulations of Fgf9 complex. (A)** Molecular dynamic simulation graph Temperature vs. Time (Production Step) **(B)** Molecular dynamic simulation graph Total energy vs. Time (Production Step)

**Figure 13.**
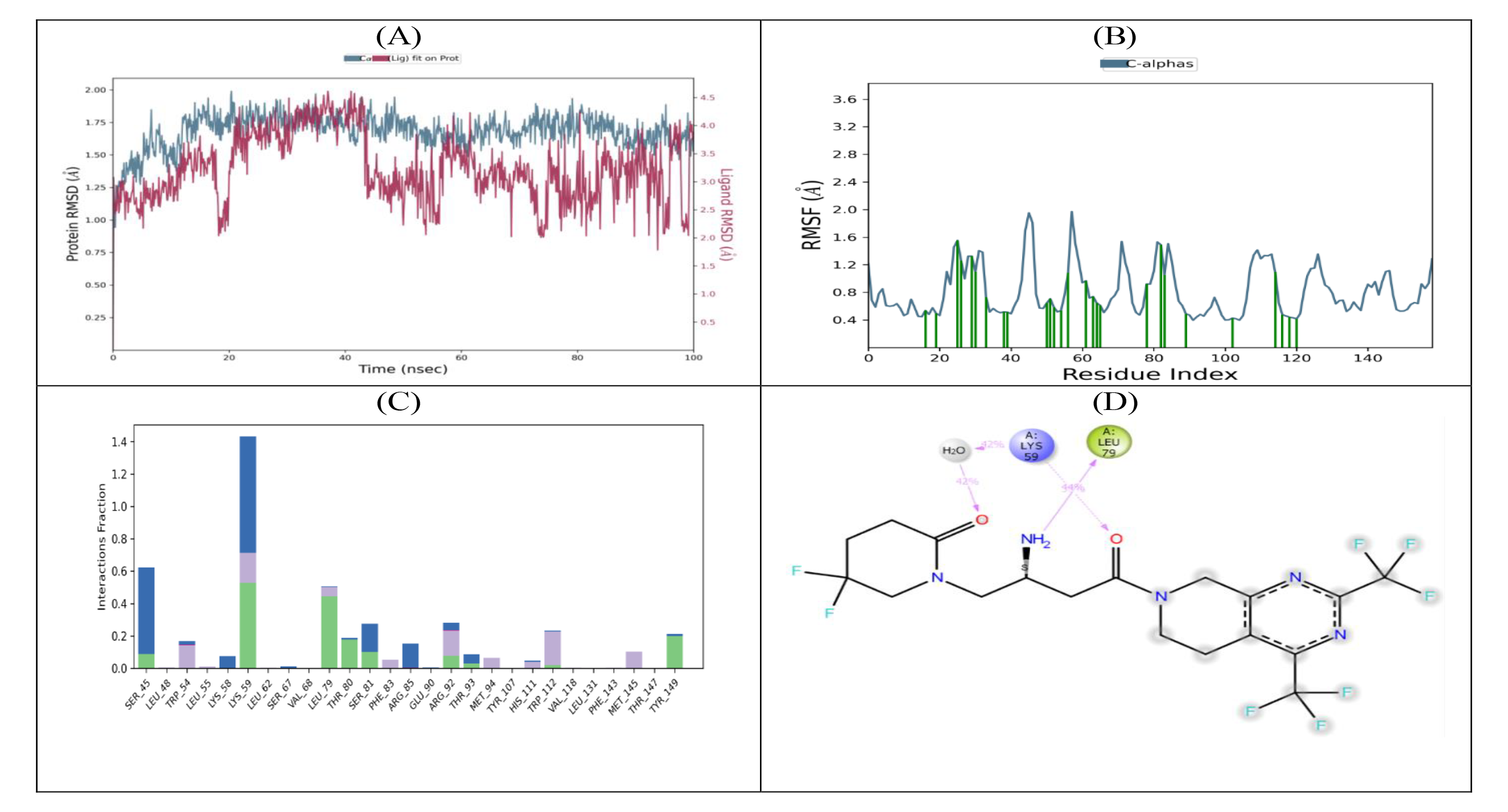
**Results of MD simulations of Ptgds complex. (A)** Molecular dynamic simulation graph Temperature vs. Time (Production Step) **(B)** Molecular dynamic simulation graph Total energy vs. Time (Production Step).

### 3.14 Molecular dynamics

For exploring nature of protein-ligand complexes at an atomistic level, molecular dynamic simulations are one of indispensable methods. Therefore, to analyse stability of ligand’s MD simulation was performed for 100ns with FGF9 and PTGDS proteins.

The RMSD analysis of FGF9 backbone revealed that after 40 ns system stabilized for best ligand. To check for flexibilities of individual residues contributing to overall fluctuations in system RMSF was computed for all protein-ligand complexes. Interaction analysis based on molecular dynamics revealed that ligands form hydrogen bonds withFGF9 protein backbone via Val192 and Arg 63. Hydrogen bonding is prominent with Val192 residue with connection probability of 75% (Figure12)

The RMSD analysis of PTGDS backbone revealed that after 20 ns system stabilized for best ligand. To check for flexibilities of individual residues RMSF was also computed for all protein-ligand complexes. Interaction analysis based on molecular dynamics revealed that ligands form hydrogen bonds with PTGDS protein backbone via LYS59, Leu79, THR80 and TYR149. Hydrogen bonding is prominent with LYS59 and LEU79 residue with connection probability of 44% (Figure13)

### 3.15 MMGBSA Analysis

To estimate relative free energy of ligand binding, Molecular Mechanics with Generalized Bonn Surface Area (MM-GBSA) was computed. From100 snapshots taken from last 10% of MD simulation trajectories, it was observed that binding free energy of DB12884 (−70.60 kcal mol^−1^) with 1ihkf is lesser than other compounds. Further, breakdown of energy terms reveals that contribution of coulomb energy is less, and lipophilic and van der Waal are much lower (Table 8). With same parameters, binding free energy of PTGDS and ligands are analysed. We selected DB12412 as the best ligand against4oru even though binding free energy was not the lowest as compared to other compounds but MD simulations studies revealed it as most stable compound making complex with PTGDS (Table 9).

**Table 8.** Binding free energy of FGF9 protein-ligand complexes calculated using MM/GBSA method

| Ligand | Binding Free Energy | Coulomb Energy | Lipophilic Energy | van der Waals energy |
| --- | --- | --- | --- | --- |
| 1ihkF +(DB08019) | -64.41174641 | -12.25871472 | -25.19157853 | 0.276049003 |
| 1ihkF +(DB09350) | -59.05395971 | -9.948418899 | -21.24204441 | -43.32924185 |
| 1ihkF+(DB12313) | -63.93803761 | -15.85440901 | -26.3480016 | -42.75472744 |
| 1ihkF +(DB12884) | -70.60573981 | -15.29025998 | -24.00579309 | -53.42542006 |
| 1ihkF +(DB08018) | -68.48405337 | -15.05676304 | -26.81736591 | -50.30653008 |
| 1ihkF +(DB06506) | -62.60080508 | -2.500560445 | -22.2636050 | -49.10056532 |
| 1ihkF +(DB06106) | -64.70404662 | -63.52808327 | -23.71529659 | -63.47729387 |
| 1ihkF +(DB13048) | -64.70404662 | -63.52511775 | -23.71529659 | -63.51116715 |

**Table 9.** Binding free energy of PTGDS protein-ligand complexes calculated using MM/GBSA method

| Ligand | Binding Free Energy | Coulomb Energy | Lipophilic Energy | van der Waals energy |
| --- | --- | --- | --- | --- |
| 4oru+(DB07548) | -74.807838 | -13.68 | -27.8272 | -57.7847 |
| 4oru+(DB08749) | -67.5374 | -10.8125 | -28.596 | -60.3277 |
| 4oru+(DB12412) | -42.9621 | -6.6948 | -12.8547 | -42.3635 |
| 4oru+(DB12523) | -60.61066 | -16.5625 | -16.402 | -53.7057 |
| 4oru+(DB04957) | -69.2628 | -8.2679 | -20.5839 | -57.0595 |

## 4. Discussion

Despite enormous studies conducted for understanding sex determination and gonadal differentiation, there are still links missing that provide clues for identification of potential regulators involved in these mechanisms along with their related disorders. Different parameters have been explored using various approaches to fill gaps in knowledge. This is first *in silico* study to investigate PPI network of pathways in mammalian sex determination to explore the contributions of proteins coding and non-coding genes associated with gonadogenesis using different features and parameters. PPIN provide vast information about regulatory aspects of biological pathways and protein functions within cell which will assist in drug target identification. Investigations and topological analysis have shown that highly interacting proteins includes important and crucial regulators like enzymes, transcription factors, etc. It has been a long-standing goal in systems biology to find relations between topological properties and functional features of protein networks. However, most of focus in network studies has been on highly connected proteins i.e., Hub nodes. As a complementary notion, it is possible to define bottlenecks as proteins with a high betweenness i.e., network nodes that have many shortest paths connecting them, which can provide a more significant output.

### 4.1 Protein-protein interaction and molecular players of gonadogenesis

In present study, we constructed an undirected PPI network having 810 nodes and 12377 edges. The structural properties indicated that network shows a high connectivity and biologically informative. Cytohubba topologically analyzed relationship between nodes and edges by measuring centralities based on two categories, Local-based methods and global-based methods. The highly interacting nodes obtained from analysis was overlapped with our RNA-seq data to exclusively identify crucial differentially expressed genes. This include eight male-specific and six female-specific genes in local methods. In global methods nine genes are male-specific and four genes are female-specific. The key genes identified shows a large number of interactions and connect crucial pathways. Hub and bottleneck nodes in network indicate that these nodes are key mediator proteins with dynamic functional properties. This implies that key nodes are much more significant indicators of essentiality over “hub-ness” (Yu et al, 2007).topological analysis helped to refine network and revealed only highly interacting nodes. To further understand importance of network, we utilized MCODE plug-in of Cytoscape to interpret closely interconnected regions from network. Top three clusters with highest score and differential genes were preferred. Screening gene clusters could help in determining hub genes and their interaction along with association between these cluster genes and sex determining processes. To explore involvement of clusters in biological functions and pathways we conducted GO and KEGG pathway analysis. We found that these clusters

### 4.2 Functions of crucial nodes in PPI

We classified data by a variety of characteristics, identifying several subgroups of regulators consisting of potential players regulating sex determination, gonadogenesis. Gene ontology and pathway analysis revealed that key nodes are certainly involved in various functions associated to gonadogenesis. We identified many proteins that localize to nuclear compartment or cytoplasm having a potential role in gene regulation, transcript processing, and RNA metabolism and protein activity mediating multiple pathways and bioprocesses. These proteins are involved in RNA modification, gene expression, and regulation of DNA replication, protein synthesis, molecular transport, and signaling (Ewen et al, 2009). Most of differentially expressed genes are involved in basic processes of gene expression, RNA metabolic processes. Many are found to be enriched in regulation of RNA metabolic processes, chromatin silencing, brain development, reproductive processes and development processes sexual reproduction gamete generation.

### 4.3 Potential non-coding modulators of gonadogenesis

MicroRNAs are important regulators of biological processes and developmental gene expression, but their contribution to embryonic gonadal development is not well-understood. Due to role of miRNA in regulation of stability and translation of mRNA targets, we attempted to draw gene-miRNA interaction network, to analyze association between25 selected genes, which plays a crucial role in mechanism of sex determination and gonadal development, and targeting miRNA. MiRDB database was utilized and network was visualized through Cytoscape. Obtained list of miRNAs was overlapped with our smRNA-Seq data to get differentially expressed miRNAs. For male specific miRNAs, female specific targets and for female specific miRNAs, male specific targets were extracted. The twenty -six male-specific DE miRNAs were found to target thirty one DE targets obtained from11.5, 2.5 and 13.5 dpc. These included miRNAs of highly conserved let-family, few members from miRNA clusters17–92, already known for their importance in PGC development and spermatogenesis (Hayashi et al. 2008; Medeiros et al. 2011).

Our gene–miRNA interaction also highlights some well-known candidates’ in field of mouse sex determination and gonadal development. Let-7 is one of initially characterized miRNAs, identified in a study of developmental timing in *C. elegans* (Rougvie 2001). Processing of let-7 is inhibited by presence of LIN28, which binds pre-let-7 and prevents its processing by Dicer and TRBP into mature let-7. Reduced levels of mature let-7 result inexpression of Prdm1 and Lin28 and thereby PGC specification occurs (Nie et al. 2008, Viswanathan et al. 2008).miR-140-5p is expressed in growing Leydig cells and thus in testis differentiation (Rakoczy et al 2013). Interestingly, these miRNAs are also interacting with other sex-determining genes. Tang et al., 2007 reported that miRNAs let-7 and miR-30 are abundant in oocytes. Another miRNA of miR-30 family, miR-30a was found to be involved in ovarian cancer pathways. MiRNA mir-378 is found to be differentially expressed in female and targets male genes like Sox9, Xab2, ptgds. miR-378 has been shown to regulate aromatase expression and enhances oocyte growth (Xu et al., 2011).

Other form of non-coding RNA, lncRNAs make up a large part of genome, are associated with various biological functions. LncRNAs regulates corresponding genes at transcriptional and post-transcriptional level, repressing or stabilizing proteins. Its role has been studied extensively in diverse biological pathways and physio-pathological context like developmental processes, various disorders, immune responses and cancer. Depending on localization and interaction with other molecules, lncRNA modulates chromatin organizations, alter stabilization of cytoplasmic messenger RNA, interfering signaling pathways. The best example of lncRNA mediated gene repression is dosage compensation representing XIST, which is responsible for X chromosome inactivation in mammal females. This inactivation occurs when XIST molecules spread over one of X chromosomes and cause silencing of a large part of genes present on it, altogether from a different chromosome (Wutz, 2011). LncRNA, AK015184, were detected in XX but not XY germ cells from 11.5 dpc, time of sex determination and 2 days before onset of sex-specific germ cell differentiation (Chen et al. 2012).The relative position between lncRNA and its neighboring genes is a key determinant of their regulatory relationship. As widespread antisense and bidirectional lncRNA transcription was found to be evolutionarily conserved, non-random genomic distribution of lncRNAs could represent an evolutionary adaptation of genes to regulating their own expression in a context-specific manner. For instance, genomic arrangement of divergent lncRNAs is key for gene regulation in cis.

Our analysis revealed few differentially expressed lncRNA interacting with key players of sex determination pathways. We divided our selected genes based on their expression in different cell types at different time-point. Selected genes belong to bipotential gonad, some are expressing in male gonads while others only in female gonads. Emx2os-201, an antisense lncRNA gene found to interact with Cbx2 and Emx2, which are bipotential gonad genes. lncRNA shows minimum binding energy of −414.04 kcal/mol with 22 local base-pairing interaction and −1329.05 kcal/mol with 35 base-pairing interactions for Cbx2 and Emx2 respectively. Emx2os-201 shows high expression in female supporting cells that establishes gonad formation at 11.5 and 13.5 dpc. Emx2os is transcribed on strand opposite to Emx2 and overlapsEmx2 transcript displaying coordinated expression in various kind of tissues (Noonan et al. 2003).Through LncRRIsearch tool, few lncRNA was found interacting with testis-specific genes also. 2010204K13Rik-203a lncRNA located on X chromosome was found to interact with Sry transcript which is a product of Y-chromosome. The minimum binding energy and local base pairing interaction was −862.83 and 40 respectively. In database, it was showing higher expression in supporting cells of female gonad at 12.5 dpc. In MGI database, it shows slight presence in male gonads also. *Sry* initiator of male sex determination is switched off by 12.5 dpc. LncRNAs modifies chromatin arrangement during interaction of gene and lncRNA. Another, lncRNA, Gimaplos −201 also found to interact with *Sry* with minimum binding energy of −279.86 kcal/mol and 14 base pairing interaction showing expression at 12.5dpc in endothelial cells in female gonads. Other testis specific genes are also targeted by these lncRNA. Sox9 and Sox8 also interact with Emx2os lncRNA with a minimum binding energy of −1603.04 kcal/mol with 68 local base pairing and −332.8 kcal/mol with 19 base pairing respectively. As mentioned earlier that *Emx2os* is expressed in female supporting cells from 11.5 dpc onwards. Analyzing interaction data cleared that Emx2os shows great affinity for Sox9 sequences as it is having a very small sum of binding energy, indicating high interaction with gene. This could be an indication that this lncRNA might be regulating Sox9 in female cells during sex determination along with other known regulatory elements.

The study also revealed that ovary-specific genes like Wnt4 and Rspo1 are also interacting with few lncRNA. Wnt4 is found to interact with Tctn2-202 with a binding energy of −1056.07 kcal/mol and 24 local base pairing. According to Jameson database, it expresses in supporting cells of male gonads at 12.5 and 13.5 dpc. This is a newly discovered transcript with limited knowledge. This gene encodes a membrane protein, which is involved in hedgehog signaling and essential for ciliogenesis. They show varying role in HH signaling on other hand a conserved function in Gli processing and neural tube patterning (Wang et al., 2017).The lncRNA, small nucleolar RNA host gene 1 (SNHG1), located at 11q12.3,promotes progression of cancers and decreasesapoptotic activity in them. It promotes cell growth, invasion and migration in pancreatic cancer. This activation takes place through Notch-1 signaling pathway in pancreatic cancer. Down-regulation of Snhg1 inhibited activation of Notch pathway and inhibited expression of other related genes (Cui et al. 2018). As shown in data (Table 6.4) it is found to show strong interaction with WNT4 having minimum binding energy of −592.75 kcal/mol and 26 local-base pairing interactions. Expression profiling revealed that it is enriched in 11.5 & 13.5 germ cells in both male and female gonads but were depleted in male supporting cells at 12.5 dpc. Study conducted by Chen et al. 2020 revealed that Snhg1 activates Wnt/B-catenin and PI3K/Akt/mTOR signaling pathway mediated by *EzH2* gene. This synergistic action affects proliferation, autophagy and apoptosis of prostate cancer. These results strongly indicated that lncRNA Sngh1 could promote Wnt and Notch signaling which will further increase proliferation of gonadal cells thus regulating gonad differentiation.Rspo1 also interacts with Emx2os with binding energy of −510.05 kcal/mol and 20 local base pairing.

### 4.4 Computer-aided drug designing and prediction of potential candidates for combating DSDs

Variation or any defects in key genes related to gonadal development may lead to critical abnormalities. Many diseases and clinical conditions are already being reported, which include gonadal dysgenesis, ovo-testicular conditions, tumorigenesis, cyst formation, cancer, etc. For a disease-drug association and considering candidates to pursue as potential drug targets it is important to analyze certain factors, such as specific disease or condition being targeted, mechanism of action of drug, and availability of existing treatments, etc.In this study, we adopted a novel computational approach for predicting new genes related to fundamental pathway of sex determination and gonad development. This integrates a protein-protein-interaction network showing different regulatory elements and finding new drugs for a gene involved in DSDs through molecular docking. For molecular docking purpose, we analyzed all 25 genes that could provide complete 3D structure of protein. Out of all Ptgds and Fgf9 were found to have an appropriate 3D structure that is almost complete with a good duggable activity. Further analysis and validation of structure using PROCHECK module, it was revealed that in comparison to all other protein structures Ptgds (4oru) and Fgf9 (1IHK) gave a perfect Ramachandran plot with zero residues in disallowed region. In addition to enlisted importance of Fgf9, this fact made Fgf9 a suitable candidate for molecular docking and prediction of novel drugs against it. We collected 3D structure of several drugs and tested them for docking analysis.

Docking results suggested two potential inhibitorsDB12884 and DB12412 with binding energies of −38.5338 kcal/mol and −48.1254 kcal/mol respectively. Compound DB12884 targeting FGF9 is known as Lavoltidine. It belongs to investigational group in Drugbank. Currently, it is being used in trials for diagnosis of reflux and other gastroesophageal diseases. Chemically it is a trizole and a highly potent antagonist for H2 receptor. Drug is not fully annotated but few interaction known are with HRH2, ARE and REG1. It is also known to interact with many drugs, one of them is Fluconazole, which is a well-used antifungal drug in market. Other drug, Gemigliptin which shows best interaction with PTGDS is a dipetidyl peptidase inhibitor used in type 2 diabetes and cancer. It is also under investigational group of FDA approved drugs which is being used in combination with other diseases for diabetes and treatment of cancer. Some drugs that are interacting with this are Leuprolide, Goserelin, Octreotide, etc. These drugs are gonadotropin releasing hormone agonist and an anti-estrogen.

## 5. Conclusion

We present here the first large scale bioinformatics survey of phenomenal mechanism of mammalian sex determination and gonadogenesis to provide a valuable resource for understanding molecular panorama of underlying mechanism. Considering popularity of performing bioinformatics analysis, we designed this work in study of gonadogenesis and related DSDs. A comparison of differentially expressed genes/proteins either between sexes or at different developmental stages establishes a preliminary basis of further studies. Along with prediction of putative genes involved in sex determination and sexual development, we analyzed regulatory mechanisms also. Although progress has been done in understanding DSDs, a dearth of knowledge is still surrounding. With the existing findings, we predicted compounds interacting with already known elements of sexual development, which will potentially regulate specific protein, combating disorders of sexual development.

## Acknowledgments

We thank Prof R Raman of our lab for initiating research on sex determination. We acknowledge CSIR, New Delhi for providing fellowship to CK; UGC & DST for strengthening research facilities in department through UGC-CAS, DST-FIST, and IoE programs.

Computer system and online applications of School of Biotechnology, BHU was used for most of the work. The simulations were performed on resources provided by ‘PARAM Shivay Facility’ under the National Supercomputing Mission, IIT-BHU, Varanasi are gratefully acknowledged.

## Author Contributions

The concept and design of study was designed by CK. *In silico* analyses was completed by CK and VKS. Analyses of data is done by CK and VKS. CK wrote the first draft of manuscript. JKR and VKS helped in writing. All authors have read and agreed to the published version of the manuscript

## Declaration

No conflict of interest.

